# Spleen Tyrosine Kinase (SYK) negatively regulates ITAM-mediated human NK cell signaling and CD19-CAR NK cell efficacy

**DOI:** 10.1101/2024.07.09.602676

**Authors:** Alberto J. Millan, Vincent Allain, Indrani Nayak, Oscar A. Aguilar, Janice S. Arakawa-Hoyt, Gabriella Ureno, Allison Grace Rothrock, Avishai Shemesh, Justin Eyquem, Jayajit Das, Lewis L. Lanier

## Abstract

NK cells express activating receptors that signal through ITAM-bearing adapter proteins. The phosphorylation of each ITAM creates binding sites for SYK and ZAP70 protein tyrosine kinases to propagate downstream signaling including the induction of Ca^2+^ influx. While all immature and mature human NK cells co-express SYK and ZAP70, clonally driven memory or adaptive NK cells can methylate *SYK* genes and signaling is mediated exclusively using ZAP70. Here, we examined the role of SYK and ZAP70 in a clonal human NK cell line KHYG1 by CRISPR-based deletion using a combination of experiments and mechanistic computational modeling. Elimination of *SYK* resulted in more robust Ca^++^ influx after cross-linking of the CD16 and NKp30 receptors and enhanced phosphorylation of downstream proteins, whereas *ZAP70* deletion diminished these responses. By contrast, *ZAP70* depletion increased proliferation of the NK cells. As immature T cells express both SYK and ZAP70 but mature T cells often express only ZAP70, we transduced the human Jurkat cell line with SYK and found that expression of SYK increased proliferation but diminished TCR-induced Ca^2+^ flux and activation. We performed transcriptional analysis of the matched sets of variant Jurkat and KHYG1 cells and observed profound alterations caused by SYK expression. As depletion of *SYK* in NK cells increased their activation, primary human NK cells were transduced with a CD19-targeting CAR and were CRISPR edited to ablate *SYK* or *ZAP70*. Deletion of *SYK* resulted in more robust cytotoxic activity and cytokine production, providing a new therapeutic strategy of NK cell engineering for cancer immunotherapy.

## Introduction

Natural Killer cells recognize and directly target cancer cells without the need for prior exposure. Unlike T cells, which require the presentation of specific antigens by MHC molecules, NK cells detect and eliminate cells that display stress-induced ligands, downregulation of MHC class I, or other markers associate with cellular transformation. NK cells use of germline-encoded receptor recognition of conserved ligands is advantageous in the context of cancer where the heterogeneity of tumors makes it challenging to identify specific antigens that are universally expressed across different types of cancers. Chimeric antigen receptors (CAR), whether introduced into NK or T cells, share the same fundamental concept of being engineered to express synthetic receptors that recognize specific antigens on target cells and thus redirect killing. Development of CAR-NK cells is motivated by the unique characteristics of NK cells, such as their innate recognition abilities, broad specificity, and potential for reduced off-tumor toxicity (1).

NK cells provide rapid immune responses against viral infections and cancer. In addition to germline-encoded receptor recognition by NK cells, antibody-dependent cellular cytotoxicity (ADCC) is a mechanism that uses activating Fc receptors (FcγRIIIa, CD16a, *FCGR3A*) to provide exquisite antigen specificity. These receptors bind the Fc region of IgG and when bound to the surface of target cells can activate and release cytotoxic granules containing granzymes and perforin and secrete cytokines and chemokines (2). Human NK cells that express CD16 are identified as the mature (CD56^dim^CD16^+^) subset, whereas the lack of CD16 is a characteristic of the immature (CD56^bright^CD16^−^) NK cell subset. Memory or adaptive NK cells have been identified in individuals exposed to human cytomegalovirus (CMV) and they possess the ability to undergo clonal expansion, generate long-lasting memory responses, and display enhanced effector functions including ADCC. In humans, these adaptive NK cells are identified by the surface expression of NKG2C and often the loss of expression of FcεR1γ, SYK, and/or EAT2 (3–6). Understanding the adaptive features of NK cells, particularly in the context of ADCC (7), has broad implications for cancer immunotherapy.

ITAM signaling in NK cells provides activation through the CD16, NKp46, and NKp30 receptors and their associated adaptors FcεR1γ and CD3ζ, which form homo- or hetero-dimeric structures (e.g., FcεR1γ-FcεR1γ, FcεR1γ-CD3ζ, or CD3ζ-CD3ζ) (2). The NKG2C-CD94 complex is expressed on adaptive NK cells and non-covalently associates with DAP12 homodimers, which each carry one ITAM (8, 9). In a similar manner, T cells use the TCR complex with the associated CD3 adapters (CD3γ, CD3δ, CD3ε, and CD3ζ) (10). Upon ligand binding, these receptors activate Src-family kinases (SFKs) such as Lck that promote the phosphorylation of ITAM residues. Phosphorylated ITAMs serve as docking sites for SYK and ZAP70 kinases, which possess tandem Src homology 2 (SH2) domains that specifically bind to these phosphorylated motifs. Once bound to the phosphorylated ITAMs, SYK and ZAP70 trigger pathways involving phospholipase C-γ (PLC-γ) leading to calcium release, PI3K (phosphoinositide 3-kinase) leading to AKT activation, and Vav1 leading to ERK activation (11). Together, SYK and ZAP70 play crucial roles in transducing signals from ITAM-containing receptors that orchestrate NK and T cell immune responses.

SYK and ZAP70 belong to the family of non-receptor tyrosine kinases and transmit signals downstream of immunoreceptors (11, 12). Both kinases share similar structures such as tandem SH2 domains at their N-terminus and a C-terminal kinase domain. SFKs phosphorylate tyrosine residues within their activation loop and cause a conformational change; however, only SYK can undergo autophosphorylation without the need of SFKs (11). Furthermore, SYK and ZAP70 are differentially expressed in immune cell subsets. ZAP70 is predominately expressed in T cells, whereas SYK is expressed in B cells, mast cells, macrophages, dendritic cells, monocytes, and some γδ T cells (11). Uniquely, most NK cells co-express both SYK and ZAP70 but can downregulate *SYK* expression by methylation in adaptive NK cells (3, 4). Recent studies by Dahlvang and colleagues (13) have reported that CRISPR-mediated ablation of *SYK* in primary human NK cells increases their cytolytic activity and cytokine production, whereas ablation of *ZAP70* impairs these effector functions.

FcεR1γ and/or CD3ζ transduce signals via SYK and/or ZAP70, which makes studying these pathways and signaling molecules challenging. In this study, we have investigated the unique aspects of ITAM-induced activation by SYK and ZAP70 in human NK and T cells using a combination of experiments and mechanistic computational modeling.

## Materials and Methods

### Human samples

Primary human NK cells were obtained from healthy donors with informed consent in accordance with approval by the University of California Committee on Human Research IRB #10-00265. PBMC were obtained by Trima Residuals from apheresis collection of blood enriched for leukocytes, predominately mononuclear cells (Vitalant, https://vitalant.org/Home.aspx).

### Cell lines

The human KHYG1 parental NK cell line was generously provided by Dr. Nicolai Wagtmann (Dragonfly Therapeutics, Inc.). The human Jurkat leukemic cell line was generously provided by Dr. Arthur Weiss (University of California San Francisco, CA). HEK293T cells were obtained from American Type Culture Collection (ATCC). The SUPB15 B lymphoblast cell line was obtained from the ATCC, transduced to express mKate2 and luciferase, and cultured in IMDM with 100 U/ml penicillin, 100 µg/ml streptomycin, 50 µM 2-mercaptoethanol, and 20% FCS. KHYG1, Jurkat, and HEK293T cells were cultured in complete RPMI-1640 (RPMI) or DMEM media supplemented with 2 mM glutamine, 100 U/ml penicillin, 100 µg/ml streptomycin, 50 µg/ml gentamicin, 110 µg/ml sodium pyruvate, 50 µM 2-mercaptoethanol, 10 mM HEPES, and 10% FCS. KHYG1 cells were supplemented with 200 U/ml recombinant human IL-2 (Teceleukin; provided by NCI Biological Resources Branch). Primary human NK cells were cultured in NK MACS medium (Miltenyi) supplemented with 5% human platelet lysate (EliteGro-Adv, Elite Cell), 0.5% penicillin-streptomycin (Gibco), and recombinant human IL-2 1000 U/mL (Peprotech).

### Antibodies and flow cytometry

Fluorochrome-conjugated antibodies were used to stain surface and intracellular proteins. Cells were first stained at 4° C in Cell Staining Buffer (BioLegend) for 30 min, washed, and analyzed using a LSRII (BD Biosciences) or spectral Cytek Aurora (Cytek Biosciences). Intracellular staining required additional steps after surface staining using the BD Cytofix/Cytoperm^TM^ Fixation/Permeabilization Solution Kit (BD Biosciences) following the manufacturer’s protocol. FcR were blocked using human TruStain FcXTM (BioLegend). Zombie Red (BioLegend) was used for Live/Dead exclusion dye. Data were analyzed using FlowJo software (FlowJo) by gating on lymphocytes identified based on forward and side light scatter properties and by excluding doublets, dead cells, and lineage-negative (CD14-CD19-) cells. Anti-NKp30 (clone P30-15), anti-CD56 (clone NCAM16.2), anti-NKp46 (clone 9E2), anti-CD3 (clone UCHT1 and OKT3), anti-CD16 (clone 3G8), anti-CD57 (clone QA17A04), anti-Fc R1antibody subunit (product # FCABS400F), anti-CD45 (clone 2D1), anti-SYK (clone 4D10.2), anti-PLZF (clone R17-809), anti-NKG2A (clone REA1100), anti-NKG2C (clone 134591), anti-ZAP70 (clone A151148 or 1E7.2), anti-CD19 (clone H1B19), anti-CD14 (clone 63D3), anti-CD69 (FN50), anti-CD247 (clone 6B10.2), anti-CD5 (clone UCHT2), anti-G4S Linker for CAR detection (clone E7O2V), anti-CD107a (clone H4A3), anti-TNF (Mab11), and anti-IFN (clone W19227A) were purchased from BioLegend, BD Biosciences, Cell Signaling, Milli-Mark, or ThermoFisher.

### Genomic CRISPR-Cas9-based editing

CRISPR Cas9-RNP single guides were used to delete endogenous human *SYK* and *ZAP70* genes from KHYG1 cells. CRISPR single guide sequence (5’-ACACCACTACACCATCGAGC-3’; IDT) was used to target exon 2 of human *SYK* and sequence (5’-TCATACACGCTCGTGTCCAT-3’; IDT) was used to target exon 2 of human *ZAP70*. In brief, KHYG1 cells were electroporated (4D-Nucleofector^TM^ System; Lonza) using the P3 Primary Cell Nucleofector Solution (Lonza), 120 bp ssDNA non-targeting electroporation stabilizer (IDT), and the corresponding CRISPR target guide RNA. KHYG1 cells were then single-cell sorted into 96-well plates and grown in RPMI-1640 media supplemented with recombinant human IL-2 1000 U/mL (Peprotech) for 5-6 weeks. Intracellular flow cytometry was then used to screen for single-cell clones of either *SYK*- or *ZAP70*-depleted KHYG1s cells. Four *SYK*- and four *ZAP70*-ablated single cell clones were combined and used. Depletion of endogenous *SYK* or *ZAP70* was confirm using intracellular flow cytometry and Western blot analysis.

### Retroviral transfection and transduction

Human SYK or CD16 cDNA sequences were cloned into pMXs-Puro retroviral vectors. Retrovirus were generated using HEK293T cells plated 1-day prior in 6-well plates and transfected using the Lipofectamine 2000 reagent according to the manufacturer’s protocol (ThermoFisher Scientific), pMXs-Puro vector, and packaging plasmids. HEK293T viral supernatant was collected and used to transduced KHYG1 or Jurkat cells. Surface expression of CD16 was used to sort CD16-expressing KHYG1 cells. Jurkat cells were transduced with pMXs-Puro-SYK and then selected with puromycin for 5-6 weeks. Expression of SYK in Jurkat cells was determined using intracellular flow cytometry with purity of >95% and ZAP70 expression matched that of the parental cells.

### Western blotting

Cell protein was prepared using RIPA Lysis Buffer (ThermoFisher Scientific) following the manufacturer’s protocol. Protein samples were first reduced using 355 mM 2-mercaptoethanol (Bio-Rad) in Laemmli Sample Buffer (Bio-Rad). Proteins were separated on 12% Mini-PROTEAN^TM^ TGX^TM^ precast protein gels (Bio-Rad) and then blotted to PVDF membranes using a 20% methanol pre-wet type blotting system and Criterion Blotter with Plate Electrodes (Bio-Rad). Membranes were washed with TBST (Bio-Rad) and incubated overnight at 4° C or for 2 hours at room temperature with blocking buffer (5% skim milk in TBST). The blots were washed and probed with 0.5-1 ug/ml of primary antibody against SYK (clone 4D10.2; BioLegend) or ZAP70 (clone 1E7.2; BioLegend) for 1-2 hours with gentle rocking. Protein loading control HRP was detected using anti-GAPDH (clone W17079A; BioLegend). The blots were then washed 3 times with TBST and incubated with secondary antibody, horseradish peroxidase-conjugated goat anti-mouse IgG (BioLegend), for 1 hour at room temperature. Membranes were incubated in Chemiluminescent substrate (SuperSignal^TM^ West Pico Plus; ThermoFisher Scientific) for 5 min and visualized using a ChemiDoc Imaging System (Bio-Rad).

### Phosphorylation kinetics assay

Cells were stimulated by antibody-induced receptor cross-linking, probed with multiple phosphorylated antibodies, and analyzed by flow cytometry. Cell numbers were first normalized between groups by using the Countess^TM^ 3 FL Automated Cell Counter (ThermoFisher Scientific). One to two million cells were stained with 5 ug/ml of biotinylated anti-CD16 (clone 3G8; BioLegend), anti-CD3 (clone UCHT1; BioLegend), or anti-NKp30 (clone P30-15; BioLegend), and Zombie Red^TM^ Fixable Viability dye (BioLegend) depending on the cell type or activation and incubated for 30 min at 4° C in HBSS (ThermoFisher Scientific) Stimulation media (HBSS with Ca^2+^ and Mg^2+^, no phenol red, 10 mM HEPES, and 1% FCS). The cells were washed and rested at 37° C for 60 min. Receptor were then cross-linked by the addition of streptavidin (500 ug/ml; BioLegend) and quenched at the conclusion of the indicated time points. The stimulation was quenched with the addition of Fixation buffer (BD Biosciences) and incubated for an additional 15 min at 37° C. The cells were permeabilized using the True-Phos^TM^ Perm Buffer (BioLegend) according to the manufacturer’s protocol. Cells were stained with anti-LYN (Tyr397)/LCK (Tyr394)/HCK (Tyr411)/BLK (Tyr389) (clone E5L3D, Cell Signaling), anti-LAT (pY226) (clone A20005, BioLegend), anti-ZAP70 (Tyr493) (clone A16043E, BioLegend), anti-ZAP70 (Tyr292) (clone A16038B, BioLegend), anti-MAPK (Thr180/Tyr182) (clone A16016A, BioLegend), anti-ERK1/2 (Thr202/Tyr204) (clone 6B8B69; BioLegend), anti-AKT (pS473) (clone M89-61, BD Bioscience), anti-SLP-76 (Ser376) (clone E3G9U, Cell Signaling), and anti-PLCγ1 (Tyr783) (clone A17025A, BioLegend) for 30 min at room temperature. The cells were then analyzed on the spectral Cytek Aurora analyzer (Cytek Biosciences).

### Calcium influx assays

T and NK cells were labeled with 7 ug/ml of Indo-1 AM (ThermoFisher Scientific) for 30 min, washed, and stained for an additional 30 min at 4° C either with biotinylated anti-CD16 (clone 3G8, BioLegend) or anti-CD3 (clone UCHT1, BioLegend). Cells were then stimulated by cross-linking the receptors with mAb and streptavidin in a manner similar to phosphorylation kinetic assays (BioLegend). Cells were analyzed on the flow cytometer with UV fluorescence channel detectors by acquiring the first 30 seconds of baseline calcium levels and then measuring calcium influx levels upon the addition of streptavidin (500 ug/ml, BioLegend) while maintaining 37° C temperature using a water pump warmer.

### Primary CAR-NK cell engineering

The cell engineering platform for expansion and genome editing of primary human NK cells were adapted from Huang *et al.* (14). Primary human NK cells were isolated from PBMC obtained after density gradient centrifugation of Trima Residuals from apheresis collection (Vitalant). Negative selection of NK cells was performed by using a EasySep™ Human NK Cell Enrichment Kit (Stemcell). Cells were cultured in NK MACS medium (Miltenyi) supplemented with recombinant human IL-2 1000 U/mL (Peprotech). Cells were activated over 7 days with anti-CD2 and anti-NKp46-coated beads (Miltenyi) and cultured at 1 × 10^6^ cells/mL. At day 7, beads were removed, and NK cells were transduced by centrifugation on Retronectin (Takara)-coated plates with a concentrated (Retro-X™ concentrator, Takara) SFG γ-retroviral vector encoding a second generation anti-CD19 Chimeric Antigen Receptor (CAR) with CD28 co-stimulation and CD3ζ domains packaged using 293Vec-RD114 cells (Biovec Pharma). Genes of interest were subsequently deleted by CRISPR editing in transduced NK cells. After 2 days of rest in complete NK cell medium, 2 × 10^6^ NK cells were electroporated with 40 pmol of purified Cas9-NLS protein (MacroLab) complexed with 80 pmol of sgRNA (Synthego) using a 4D nucleofector (Lonza) in 20 μl of P3 buffer and supplement (Lonza) using the CM-137 pulse code. Immediately after electroporation, edited NK cells were rescued in prewarmed culture medium and plated for further cell culture.

### Flow cytometry-based 4-hour cytotoxicity assay

The cytotoxicity of NK cells in a 4-hour assay was measured by flow cytometry as adapted from Aguilar *et al*. (15). In brief, CD19-expressing SUP-B15 (ATCC) cells were labeled with CellTrace^TM^ Violet (ThermoFisher Scientific) and co-cultured with primary CAR-NK cells at varying E:T ratios. Cells were cultured for 4-hours and then washed and stained with Zombie Red^TM^ Fixable Viability dye (BioLegend). CountBright Absolute Counting Beads (ThermoFisher Scientific) were added to each well to obtain cell counts according to the manufacturer’s protocol. Percentage of specific target cell lysis was calculated with the following formula, which determines the number of live cells in experimental samples relative to control samples:

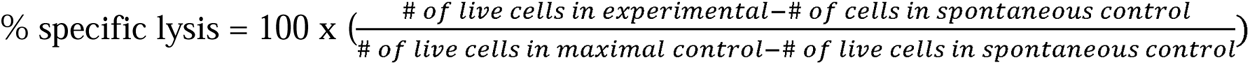

### Luciferase-based 24-hour cytotoxicity assay

The cytotoxicity of NK cells was determined by using a standard luciferase-based assay. In brief, SUP-B15 cells were transduced with lentivirus to express firefly luciferase-mKate2 and served as target cells. The effector (E) and tumor target (T) cells were co-cultured in triplicates at the indicated E:T ratios using white-walled 96-well flat clear-bottom plates with 5 × 10^4^ target cells in a total volume of 100 μL per well in NK cell medium without IL-2. The control for maximal signal was SUP-B15 cells alone, and the control for minimum signal was SUP-B15 cells with Tween-20 (0.2%). Co-cultures were incubated for approximately 20-24 hours. Then, 100 μL D-luciferin (GoldBio, 0.75 mg/ml) was added to each well, and luminescent signal was measured using a GloMAX Explorer microplate reader (Promega).

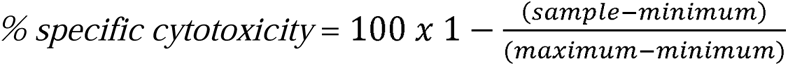

### Next-generation sequencing

Total RNA was extracted from cells using the Qiagen RNeasy Mini kit with DNase treatment per the manufacturer’s protocol. Illumina libraries were generated from total RNA using the Universal Plus mRNA-Seq with NuQuant kit and sequenced on the HiSeq to a depth of ≥66 million reads. Sequences were align to the Ensembl human genome build GRCh38 annotation version 95 using STAR version 2.7.5c (16). Differentially expressed genes were analyzed between groups using DESeq2 version 1.42.1 (17) on R software version 4.3.3.

### Graph statistical analysis

Data were analyzed using Prism version 10.2.3 (GraphPad) by unpaired and paired t-test. Graphs show mean standard deviation. *, P <0.05; **, P <0.01; ***, P <0.001. Additional statistical analysis details can be found in the figure legends.

### In silico modeling membrane proximal CD16 signaling

We developed a rule-based multi-compartment model describing CD16 receptor-initiated membrane proximal biochemical signaling reactions where the participating molecules are well-mixed (18–20). The model is composed of CD16 receptor, anti-CD16 antibody, CD3ζ adaptor, SFK Lck, SYK-family kinases ZAP70 and SYK, phosphatase SHP-1, and ubiquitin ligase Cbl (21–25). The signaling protein species are placed within a two compartmental simulation box representing thin layers of the extracellular region (compartment 1) of depth 2 nm and the cytosolic region (compartment 2) of depth 1 μm partitioned by the two-dimensional NK cell plasma membrane of area 5 μm × 5 μm. Plasma membrane bound CD16 binds with the Fc region of the antibody ligands in compartment 1, while the transmembrane domain of CD16 binds with various signaling molecules in compartment 2. CD16, Lck, and CD3ζ adaptors are always anchored to the membrane. ZAP70, SYK, and SHP-1 are cytosolic molecules that bind to the ITAM present on the tails of the CD3ζ proteins extended in the cytosol. Cbl, a cytosolic ubiquitin ligase, binds to the ITAM through ZAP70 or SYK phosphorylation (26, 27). The major signaling reactions in the simulation box are shown in Fig. 2G and discussed in our GitHub link. The details of the signaling reactions, the associated rates, and abundances (or copy numbers) of CD16, antibody ligands, and signaling proteins used in the model are shown in Table S1. Each ITAM on CD3ζ can exist in ten distinct states including fully and partially unphosphorylated state, fully phosphorylated state, fully phosphorylated state bound to ZAP70 (28), fully or partially phosphorylated states bound to SYK and/or SHP-1, respectively (29, 30). Because homodimers of CD3ζ adaptor have 6 ITAMs, it can generate up to ∼6^10^ possible intermediate complexes in the model and increases computational runtime significantly when ordinary differential equations (ODEs) describing mass action rate kinetics are used. To save computational cost, we simulated the biochemical signaling reaction kinetics using stochastic Network Free Simulator (NFSIM) application (20, 31). The NFSIM application simulates the kinetics with intrinsic noise fluctuations using Gillespie’s algorithm (19).

### Minimal Model for Ca^2+^ signaling

We developed a minimal model for Ca^2+^ kinetics initiated by phosphorylated forms of ZAP70 and SYK in the NK cell by modifying a coarse-grained model of Ca^2+^ kinetics in *Xenopus* oocyte cells (32). In our model, the Ca^2+^ kinetics for pZAP70 or pSYK are given by two coupled nonlinear ordinary differential equations (ODEs) in terms of the concentration ([Ca^2+^]) of Ca^2+^ in the cytosol and the dimensionless variable *h*, which represents the proportion of activated (open channel) IP3 receptors (IP3-R) on the endoplasmic reticulum (ER) that release Ca^+2^ from ER. The ODEs contain parameters b, k_1_, k_2_, which are related to the rates of Ca^2+^ release through IP3-R from the ER to the cytosol in the basal (parameter b) and activated state (parameters k_1_ and k_2_) (32, 33)), and γ, which is related to the rate of Ca^2+^ pumped out of the cytosol (Table S1). In the ODEs, the average concentrations of pZAP70 [pZAP70] and pSYK [pSYK] at any time *t* regulate the production of cytosolic Ca^2+^ and the proportion of activated IP3 receptors. Since, there are differences in signaling reactions downstream of pZAP70 and pSYK that give rise to Ca^2+^ flux, we choose different values for some of the parameters for pZAP70- and pSYK-induced Ca^2+^ flux kinetics model. The ODEs for cytosolic [*Ca*^+2^]*_i_* induced by concentrations pZAP70 ([pZAP70](*t*) ≡ *y_z_* (*t*) or pSYK ([pSYK](*t*) ≡ *y_s_*) are given below. Here, subscript *i*=*Z* (or *S*) denotes stimulation by pZAP70 (or pSYK).

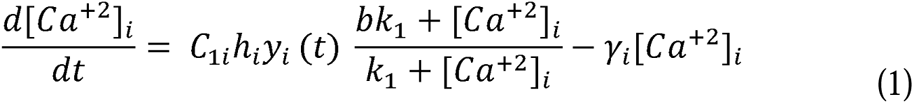

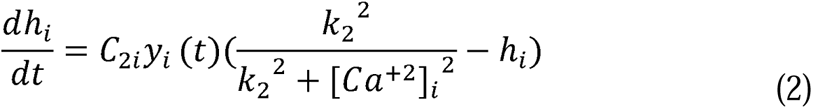

We used different parameter sets {*C*_1*Z*_, *C*_2*Z*_, γ*_Z_* } and {*C*_1*S*_, *C*_2*S*_, γ*_S_* } for pZAP70- and pSYK-induced Ca^2+^ flux kinetics model (see estimated values in Table S1). When both ZAP70 and SYK are present in the cell, the total cytosolic Ca^2+^ concentration is given by, [*Ca*^+2^]*_total_* = [*Ca*^+2^]*_Z_* + [*Ca*^+2^]*_S_* (see Fig. S1A). In experiment, Ca^2+^ flux (Fig. 2E) is measured in the unit of fluorescence ratio (Ca^2+^ =UV C (high Ca^2+^)/UV C (low Ca^2+^)), which we assumed to be proportional to the cytosolic Ca^2+^ concentration [*Ca*^+2^]*_total_*, where UV C (low Ca^2+^) represents the basal value. For simplicity, we assumed the proportional factor *f* as 1 with a unit of [μM]^−1^. *Preprocessing data for parameter estimation:* We excluded the Ca^++^ flux data for ΔSYK, ΔZAP70, and WT KHYG1 cells during the resting period (30 sec) before stimulation. Therefore, in our *in silico* analysis, we modeled the Ca^++^ flux kinetics between 30 seconds and 270 sec, as shown in Fig. 2E. The starting time, t=0, of the model simulation (Fig. 2I) thus corresponds to t=30 sec in Fig. 2E.

### Pipeline for parameter estimation

We developed a rule-based CD16 signaling model to generate phospho-ZAP70 (pZAP70) or phospho-SYK (pSYK) signal. We first created the reaction rules of CD16 signaling model using the Virtual Cell (Vcell) software and exported the signaling model in .bngl (BioNetGen) format to run in Python using PyBioNetGen (PyBNG) library (19, 34). In the .bngl file (see our GitHub link), we obtained the copy number of signaling molecules in the simulation box by multiplying the species concentrations with area or volume with given in Table S1. Similarly, we used microscopic reaction rates (in the units of seconds and molecules) for the second-order biochemical reactions occurring either in the membrane or in volume (cytosolic or extracellular). We derived these microscopic rates by scaling the second order reaction rates (in units of μm^3^ s^−1^ molecule^−1^ or μm^2^ s^−1^ molecule^−1^) in Table S1 with the volume (or area) of reactions that occurs in different compartments in the model (Table S1), namely, the extracellular volume (V_e_), the intracellular cytosolic volume (V_c_), and the membrane area (A). The simulation box represents a small transmembrane region in the NK cell, and we assumed the pZAP70 and pSYK abundances in an NK cell are generated from the averages of the copy numbers from many such simulation boxes. Thus, the average of the pZAP70 and pSYK abundances were generated from independent runs of the model. We calculated the average of the copy numbers of pZAP70 and pSYK in the simulation box (or V_c_) over a few (∼3) numbers of stochastic trajectories (Fig. 2J and Fig. S1B-C). For simplicity, we assumed all the NK cells in the experiments have similar mean copy numbers of signaling proteins and ignored the cell-cell variation in the copy numbers of the proteins. Then, we input the average pZAP70 or pSYK (in concentration μM unit) in Eqns. (1) and (2) to obtain cytosolic Ca^++^ flux averaged across the cell population. We estimated parameters (see Table S1) by fitting the model generated Ca^++^ flux with experimental Ca+^+^ flux for ΔSYK and ΔZAP70 KHYG1 cells simultaneously (Fig. 2I). For that, we minimized a combined cost function (defined as *cost function* = RSS_ΔZAP70_ + RSS_ΔSYK_), which represents the sum of individual Residuals Sum of Squares (RSS) (35) between model fit and experimental Ca^++^ flux for ΔSYK and ΔZAP70 KHYG1 cells given by RSS=Σ*_t_* (*Ca*^++^*_model_* (*t*) - *Ca*^++^*_expt_* (*t*))^2^, summed at times = 0 sec to 240 sec in the interval of 1 sec (Fig. 2I). We employed the Particle Swarm Optimization (PSO) technique for parameter estimation using the *pyswarm* library using biologically relevant parameter ranges (upper and lower bounds). For PSO execution, we choose the swarm size of particles and the number of iterations to be 40 and 20, respectively.

### Predicting Ca^++^ flux for WT KHYG1 NK cell

We considered the CD16 signaling model with ZAP70 and SYK (details of reaction rules are available in our GitHub link and Table S1) to describe CD16 signaling in WT NK cells. Then we calculated the average pZAP70 and pSYK abundances in WT NK cells using the set of parameters (Table S1) estimated from the *in silico* model. Finally, we predicted the cytosolic Ca^++^ flux assuming the Ca^++^ flux is the superposition of the Ca^++^ fluxes induced by pZAP70 and pSYK, i.e., [*Ca*^++^]*_total_* = [*Ca*^++^]*_Z_*_AP70_ + [*Ca*^++^]*_SYK_* (see Fig. S1A), where [*Ca*^++^]*_Z_*_AP70_ and [*Ca*^++^]*_SYK_* are the Ca^++^ flux stimulated by pZAP70 and pSYK, respectively. We assumed the same initial (t=0) flux of [Ca^++^] for [*Ca*^++^]*_Z_*_AP70_ and [*Ca*^++^]*_SYK_*, which is half of the experimental Ca^++^ flux value at t=0 sec (Fig. 2I).

### Parameter Estimation

We first developed rule based CD16 signaling model in the Virtual Cell software (18–20). Then exported the signaling model in BioNetGen format (.bngl) and computed the average of the stochastic kinetics of the concentrations of pZAP70 and pSYK in the CD16 signaling model using PyBioNetGen library in python (34). Then, we used the above time courses of pZAP70 or pSYK as input in the ODEs in Eqns. (1–2) and solved the ODEs numerically using *odeint* function in Scipy Integrate Python library. We set up a cost function describing the Residual Sum of squares (RSS) (35) in the ratio of the Ca^2+^ flux estimated in the model and the values measured in experiments for ΔSYK and ΔZAP70 KHYG1 NK cells (Fig. 2E). We estimated parameter values (see Table S1) by minimizing the cost function in Python using Particle Swarm Optimizer library *pyswarm* (https://pythonhosted.org/pyswarm/).

### Calculating R^2^

To show goodness of our in silico model prediction, we calculated the coefficient of determination 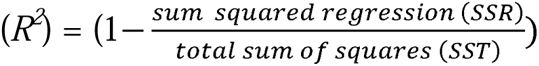

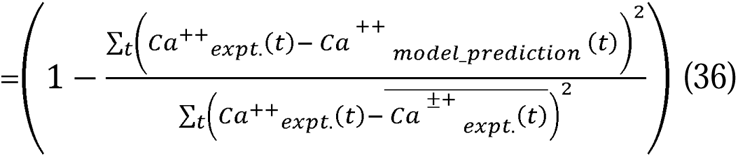

between Ca^++^ flux observed in experiments (*Ca*^++^*_expt._*(*t*)) and predicted through in silico model 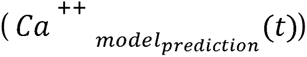 in WT KHYG1 NK cells (Fig. 2I) in time. The *t* subscript indicates the time index representing Ca^++^ flux values measured at discrete time points between 0 to 240 sec at the interval of 1 sec. 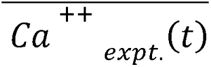 represents the average of Ca++ flux in experiments (*Ca*^++^*_expt._*(*t*)) over time.

### Details of the signaling model and availability of codes

Codes are written in python programming language. The reaction rules for Rule-based CD16 signaling model can be visualized in the Virtual cell software. The details of the signaling model and codes are available at the GitHub link: https://github.com/indraniny/ZAP_SYK_roles.

## RESULTS

### Ablation of SYK in human NK cells

Prior studies have demonstrated co-expression of SYK and ZAP70 in human NK cells. Here we determined the relative expression of these kinases in immature (CD56^bright^, CD16^−^), mature (CD56^dim^, CD16^+^), and adaptive NK cells expressing NKG2C and lacking FcεR1γ and PLZF. We analyzed 10 total donors, 5 donors (#1-5) had abundant adaptive NK cells and 5 donors (#6-10) had very few adaptive NK cells (Fig. 1A). Using 15-parameter quantitative spectral cytometry we confirmed the distinct phenotypic signature of the three major NK cell subsets found in the peripheral blood (Fig. 1B). In all donors there is progressively less SYK in the transition from immature to mature, and the lowest expression of SYK was observed in the adaptive NK cells. By contrast, ZAP70 expression increases in the immature to mature state and maintained at high levels in adaptive NK cells (Fig. 1C). To investigate the role of SYK in NK cell activation we selected a homogeneous clonal human NK cell line, KHYG1, which is typical of mature NK cells that co-expresses SYK and ZAP70. Like adaptive NK cells KHYG1 expresses CD3ζ adaptor, but not FcεRIγ. As KHYG1 cells do not express CD16 they were transduced with CD16 to evaluate ITAM-based signal transduction through the activating CD16-CD3ζ receptor. Either *SYK* or *ZAP70* were ablated by CRISRP editing technology (Fig. 2A). Complete ablation of *SYK* or *ZAP70* was confirmed by Western blot analysis and flow cytometry (Fig. 2B, C). Wildtype (WT), *SYK*-ablated (ΔSYK), and *ZAP70*-ablated (ΔZAP70) indol-1-labeled KHYG1 cells were coated with biotin-conjugated anti-CD16 and then cross-linked with streptavidin. Ca^2+^ influx was measured kinetically on a flow cytometer. Ablation of *SYK* dramatically increased the magnitude of the Ca^2+^ influx in response to CD16 stimulation, whereas ablation of *ZAP70* diminished the response compared to WT NK cells (Fig. 2D, E). In contrast to the enhanced CD16-induced Ca^2+^ response of ΔSYK KHYG1 cells, basal proliferation of ΔZAP70 KHYG1 was remarkably increased over the course of 4 days in medium with IL-2 (which is required for growth and viability of NK cells) (Fig. 2F).

**Figure 1:**
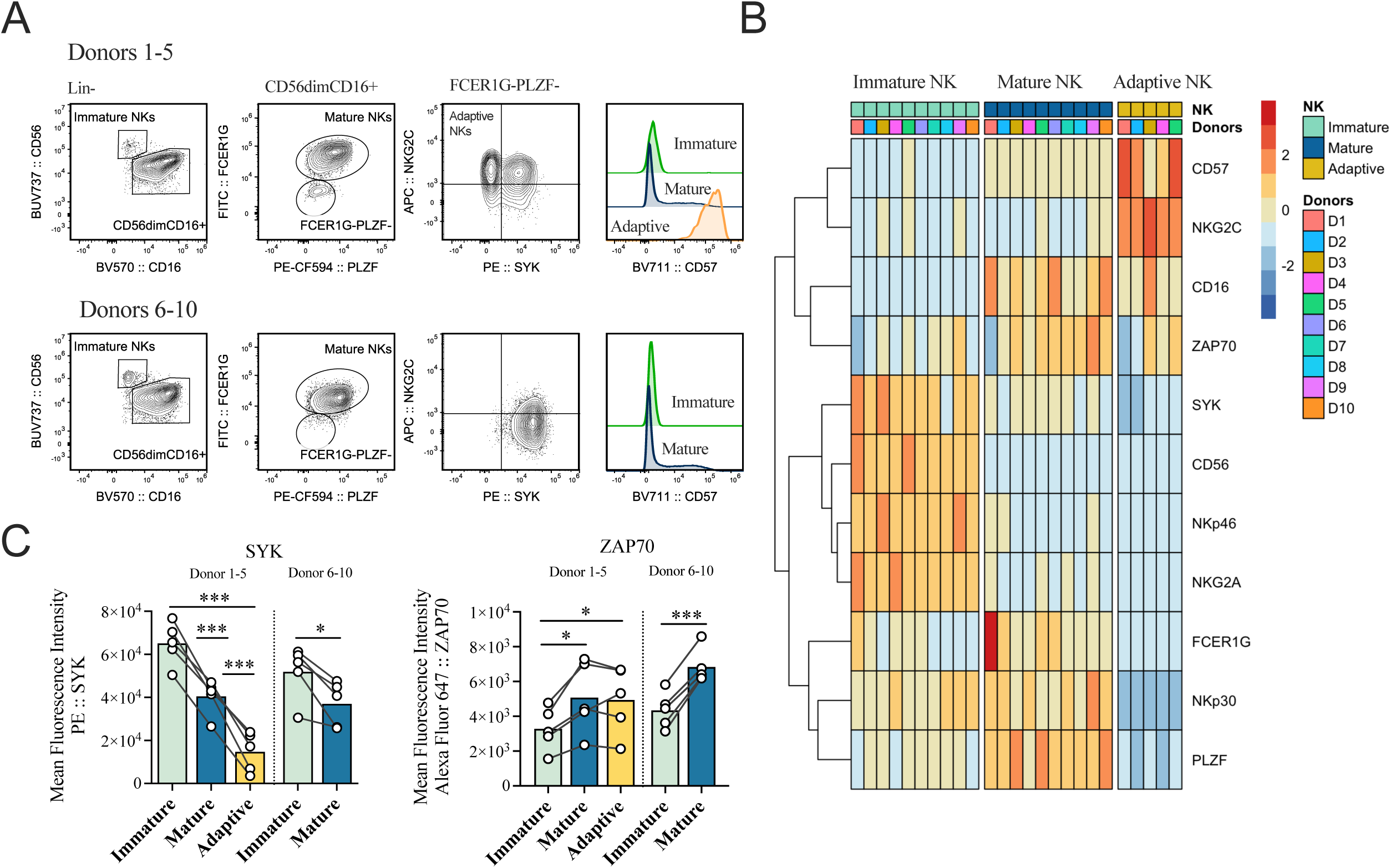
SYK and ZAP70 expression profiling in adaptive NK cell subsets. **(A-C)** Healthy human donor PBMCs were analyzed by flow cytometry for surface and intracellular markers. **(A)** Gating strategy of immature (CD56^bright^, CD16^−^), mature (CD56^dim^, CD16^+^, FcεR1γ^+^, PLZF^+^), and adaptive (CD56^dim^, CD16^+^, FcεR1γ^−^, PLZF-, NKG2C^+^) human NK cells. **(B)** Heatmap of mean fluorescent intensity (MFI) normalized by specific markers for each donor and gated on NK cell subsets. **(C)** Graphical representation of MFI data for SYK and ZAP70 for each human NK cell subset. Graphs show means with dots representing each donor; paired sample t test with significance are shown; ***, P <0.001, *, P <0.05. Total of 10 biological samples were analyzed.

We set up an *in silico* model to investigate the mechanistic roles of SYK and ZAP70 in regulating Ca^2+^ kinetics in response to CD16 stimulation in KHYG1 cells. In the *in silico* model, CD16 receptors bind to cognate ligands, and then homodimers of CD3ζ adaptors bind to the transmembrane domain of the ligand bound CD16 receptors (Fig. 2G). ITAMs associated with CD3ζ adaptors are phosphorylated by membrane bound Src Family Kinases (SFK). SYK can bind to partially and fully phosphorylated ITAMs (29, 30) (Fig. 2G right). In contrast, ZAP70 can only bind to fully phosphorylated ITAMs (28). Moreover, unlike ZAP70, SYK when bound to ITAMs can phosphorylate tyrosine residues in the ITAMs of the same CD3ζ or ITAMs of other CD3ζ proteins. The phosphorylation of ITAMs by SYK in turn can lead to increased recruitment of SYK to the ITAMs, which constitutes a positive feedback (30). ITAM-bound SYK can also auto-phosphorylate or trans-phosphorylate tyrosine residues in kinase domains on the same SYK or other SYK molecules, respectively, which increases the catalytic activity of SYK (30). The above signaling reactions involving SYK thus can give rise to positive feedback in producing catalytically active SYK molecules. Phosphorylated ZAP70 and SYK molecules are dephosphorylated by phosphatases including SHP-1 (37). In addition, the ubiquitin ligase Cbl can bind to SYK and ZAP70 complexed with CD3ζ and CD16 and degrade the entire complex (26, 27). We modeled the stimulation of intracellular Ca^2+^ flux by phosphorylated ZAP70 and SYK minimally using a coarse-grained model described by ordinary differential equations (ODEs) where the abundances (copy numbers per cell) of the phosphorylated forms of ZAP70 and SYK give rise to Ca^2+^ flux (see Materials and Methods for details). First, we trained our *in silico* model with time dependent Ca^2+^ flux in the ΔZAP70 and ΔSYK cells (Fig. 2I) and estimated several model parameters. Then, we validated our *in silico* model by predicting Ca^2+^ flux kinetics in the WT KHYG1 cells, which shows excellent agreement *(R^2^=*0.82) with the data (Fig. 2I). The model determined the following mechanism for competition between ZAP70 and SYK (Fig. 2H). The intermediate magnitude of Ca^2+^ flux in the WT KHYG1 cells compared to the lower Ca^2+^ flux in the ΔZAP70 KHYG1 cells and higher Ca^2+^ flux in ΔSYK KHYG1 cells point to a competition between ZAP70 and SYK for access of the phosphorylated ITAMs in the WT KHYG1 cells. As ZAP70 and SYK exhibit similar affinity for binding to the fully phosphorylated ITAMs with (38), the abundances of ITAM bound ZAP70 and SYK generated during early time signaling in WT KHYG1 cells are reduced compared with their counterparts in ΔSYK and ΔZAP70 KHYG1 cells (Fig. 2J). Consequently, this results in an intermediate flux of Ca^2+^ in the WT KHYG1 in between the Ca^2+^ flux observed in ΔSYK and ΔZAP70 KHYG1 cells (Fig. 2I). The Ca^2+^ flux in the WT is determined largely by pZAP70 (Fig. S1A). Furthermore, the model predicts that when the abundances of CD16 and CD3ζ adaptors are in excess compared to that of ZAP70 and SYK in the WT NK cells, the competition between ZAP70 and SYK for binding the phosphorylated ITAMs decreases (Fig. S1E-F,J-L)). The positive feedback in SYK activation along with this effect produces a greater number of pSYK and pZAP70 number in the WT compared to that of the ΔZAP70 and ΔSYK NK cells (Fig. S1H-I). Consequently, this leads to a substantially larger Ca^2+^ flux in the WT NK cells, prevailing over that of the ΔSYK and ΔZAP70 NK cells (Fig. S1D).

**Figure 2:**
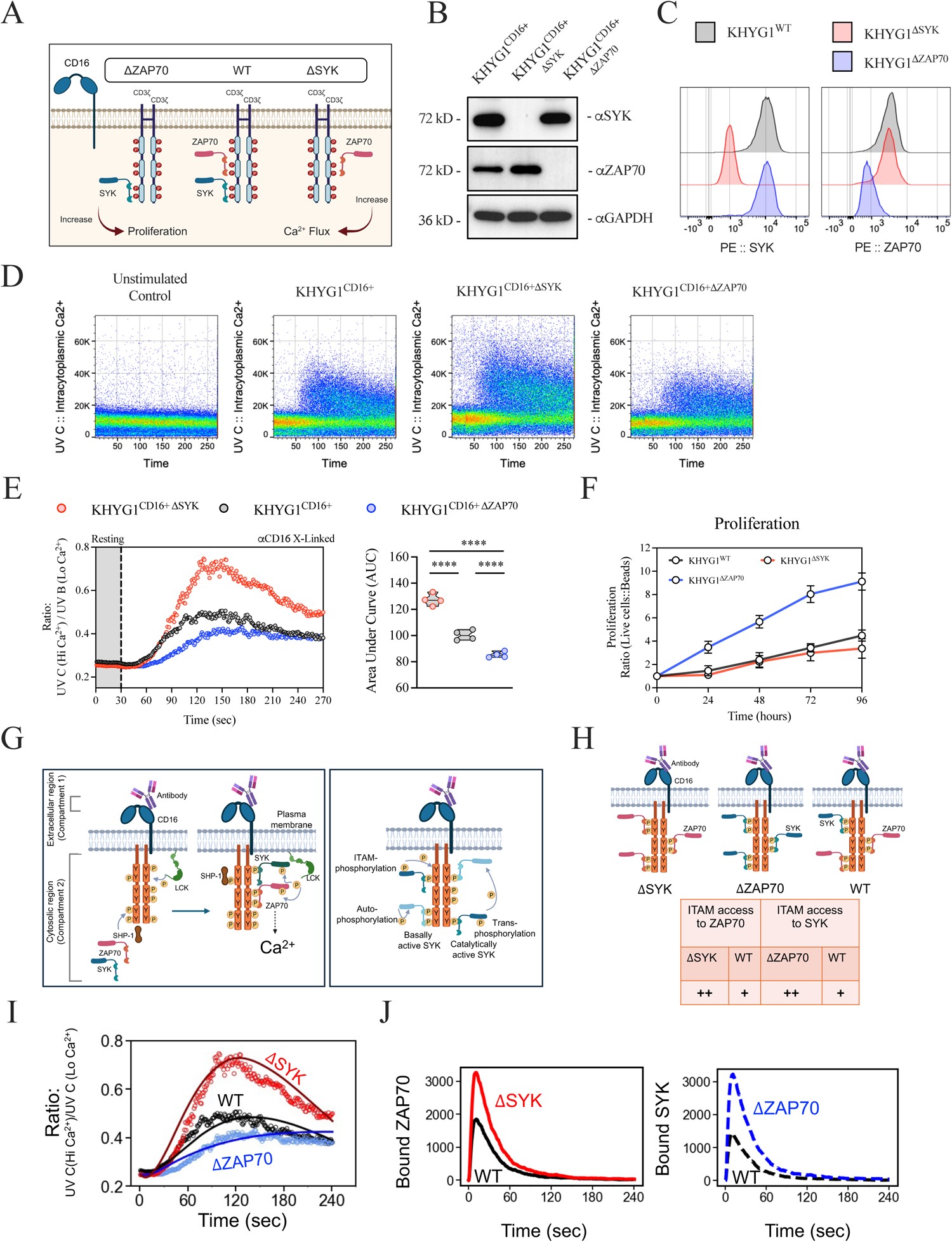
SYK and ZAP70 signaling kinetics in KHYG1 NK cells. **(A-E)** KHYG1 cells were transduced with a retroviral vector expressing human CD16 and then depleted (Δ) of either *SYK* or *ZAP70*. **(A)** Diagram of biological regulation of SYK and ZAP70. **(B-C)** *SYK* and *ZAP70* ablation was confirmed by Western blot and flow cytometric analysis. **(D)** Calcium influx assay using *SYK* and *ZAP70*-ablated KHYG1 cells. Cells were labeled with Indo-1, stained with biotinylated anti-CD16 antibody, and then cross-linked with streptavidin. Cells were collected on a flow cytometer continuously for the first 30 seconds without cross-linking and then cross-linked for an additional 270 seconds. **(E)** Graphical representative of the calcium influx for the *SYK*- and *ZAP70*-ablated KHYG1 cells; area under the curve (AUC) was calculated for each cell line, with replicates. **(F)** Cell proliferation was measured for 96 hours in media with 200 U/ml IL2. Sample t-tests with significance are shown as ****, P <0.0001. Data in D and E represent 3-4 biological replicates for two independent experiments. (**G**) (Left) Graphical representative of the major CD16 initiated signaling events involving CD16, Lck, ZAP70, SYK, and SHP-1 in the simulation box, and (Right) shows the ITAM phosphorylation by SYK, and auto-phosphorylation and trans-phosphorylation of SYK. **(H**) Graphical representative of the mechanism of increased competition between ZAP70 and SYK for accessing phosphorylated ITAMs in WT NK cells for limited number of ITAMs. (**I**) Shows model fits (solid lines) for Ca^2+^ kinetics from 0 sec-240 sec with the experimental counterparts for ΔSYK (red circles) and ΔZAP70 (blue circles) KHYG1 cells. The predicted Ca^2+^ kinetics (solid black line) for WT NK cells by the trained *in silico* model shows an excellent agreement (*R^2^* = 0.82) with the Ca^2+^ kinetics measured in WT KHYG1 cells (black circles). (**J**) Shows the number of ITAM-bound ZAP70 (left) and ITAM-bound SYK (right) in the simulation box predicted by the trained *in silico* model for WT, ΔSYK, and ΔZAP70 cells.

The kinetics in the phosphorylation of key signaling proteins downstream of ITAM-based signaling were analyzed by multi-parameter spectral cytometry (Fig. 3A, B). We observe a significant increase in the phosphorylation kinetics for SYK-ablated KHYG1 cells with a peak phosphorylation of ERK1/2 (Thr202/Tyr204) at 2 minutes when cross-linked with either anti-NKp30 or anti-CD16 (Figure 3C). Remarkably, ablation of *SYK* in KHYG1 cells stimulated through CD16 demonstrated a significant increase in many other phosphorylated proteins, including LYN (Tyr397), LCK (Tyr394), HCK (Tyr411), BLK (Tyr389), LAT (Tyr171), ZAP70 (Tyr493), MAPK (T180/Y182), ERK1/2 (Thr202/Tyr204), AKT (Ser473), and PLCy1 (Tyr783) when compared to WT KHYG1 cells (Fig. 3D, E). Phosphorylation of the tyrosine residue 292 on ZAP70 (Tyr292), which functions as a negative regulator, and SLP76 (Ser376) between *ZAP70*-ablated and SYK-ablated KHYG1 cells were unchanged (Fig. 3D, E) (39). Additionally, ZAP70*-*deficient KHYG1 cells show multiple diminished phosphorylation events compared to WT KHYG1 cells except for the combined LYN (Tyr397), LCK (Tyr394), HCK (Tyr411), BLK (Tyr389), PLCy1 (Tyr783), and MAPK (T180/Y182) (Fig. 3D, E). *SYK*-ablated cells increased many phosphorylation events of ITAM-related proteins, whereas *ZAP70*-ablation decrease phosphorylation in NK cells.

**Figure 3:**
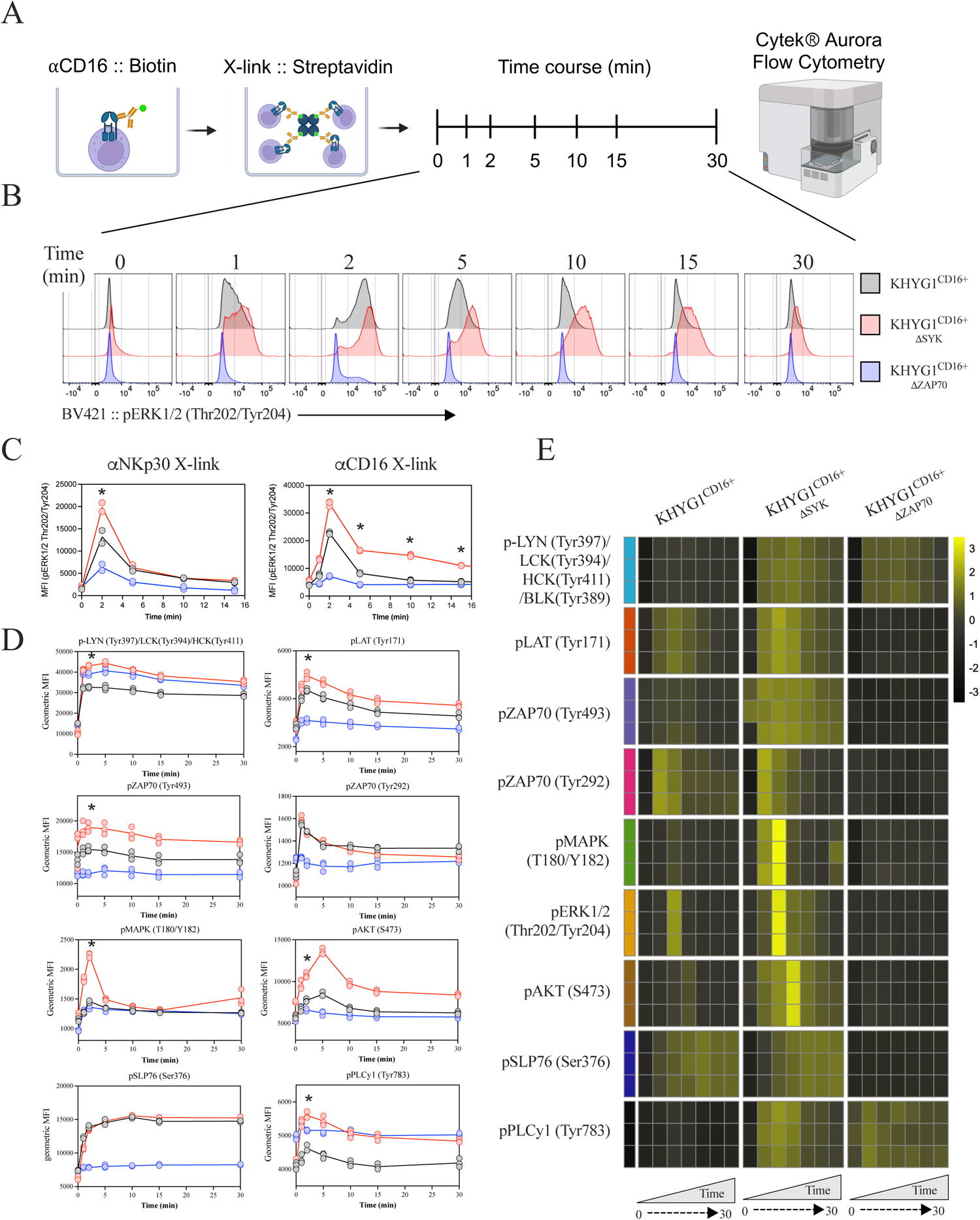
SYK depletion increases activation induced phosphorylation kinetics in KHYG1. **(A-D)** Multi-phosphorylation signaling markers were analyzed by flow cytometry for each NK cell line. KHYG1 cells were transduced with CD16 and then ablated (Δ) of *SYK* or *ZAP70*. **(A)** Diagram of KHYG1 cells stained with biotinylated anti-CD16 antibody and then cross-linked with streptavidin for 0 (unstimulated), 1, 2, 5, 10, 15, and 30 minutes and analyzed by spectral cytometry. **(B)** Representative data of phosphorylated pERK1/2 at Thr202 and Tyr204 over 30 minutes for each KHYG1 cell line. **(C)** Summarized data of pERK1/2 after cross-link activation with anti-NKp30 (left) and anti-CD16 (right). **(D)** Heatmap of geometric mean fluorescent intensity (gMFI) values overtime and normalized to rows for each phosphorylated marker. **(E)** Line plot graph of data from cells cross-linking using anti-CD16. Data used for B-D consist of 3 biological replicates.

### Effect of SYK expression on T cell signaling and proliferation

Like NK cells, immature T cells in the thymus co-express SYK and ZAP70, but SYK is downregulated as T cells mature (40), which is similar to the downregulation of SYK in human adaptive NK cells. Some subsets of mature human T cells uniquely co-express SYK and ZAP70 (41). To address the effects of SYK co-expression in a ZAP70-bearing T cell, Syk was transduced into the prototypic Jurkat T cell line, which has been used for the identification of many ITAM-based signaling pathways (Fig. 4A) (42). The expression of SYK was confirmed by Western blot and flow cytometric analysis (Fig. 4B, C). Co-expression of SYK did not affect the endogenous levels of expression for ZAP70, CD3ε, or CD3ζ or affect basal levels of CD69, indicative of no overt activation of the cells (data not shown). Anti-CD3 stimulation of the WT and SYK-bearing Jurkat cells resulted in diminished Ca^2+^ influx in the Jurkat cells expressing SYK (Fig. 4D, E) and significantly lower phosphorylation of LYN (Tyr397), LCK (Tyr394), HCK (Tyr411), BLK (Tyr389), ZAP70 (Tyr493), pZAP70 (Tyr292), and ERK1/2 (Thr202/Tyr204) compared to WT cells (Fig. 4F, G). Diminished activation in SYK-bearing Jurkat cells were also reflected by a delayed induction of CD69 after activation (Fig. 4H). Similar to the KHYG1 NK cells expressing SYK, proliferation of SYK-bearing Jurkat cells significantly increased over 4 days in culture compared to WT Jurkat cells expressing only ZAP70 (Fig. 4I).

**Figure 4:**
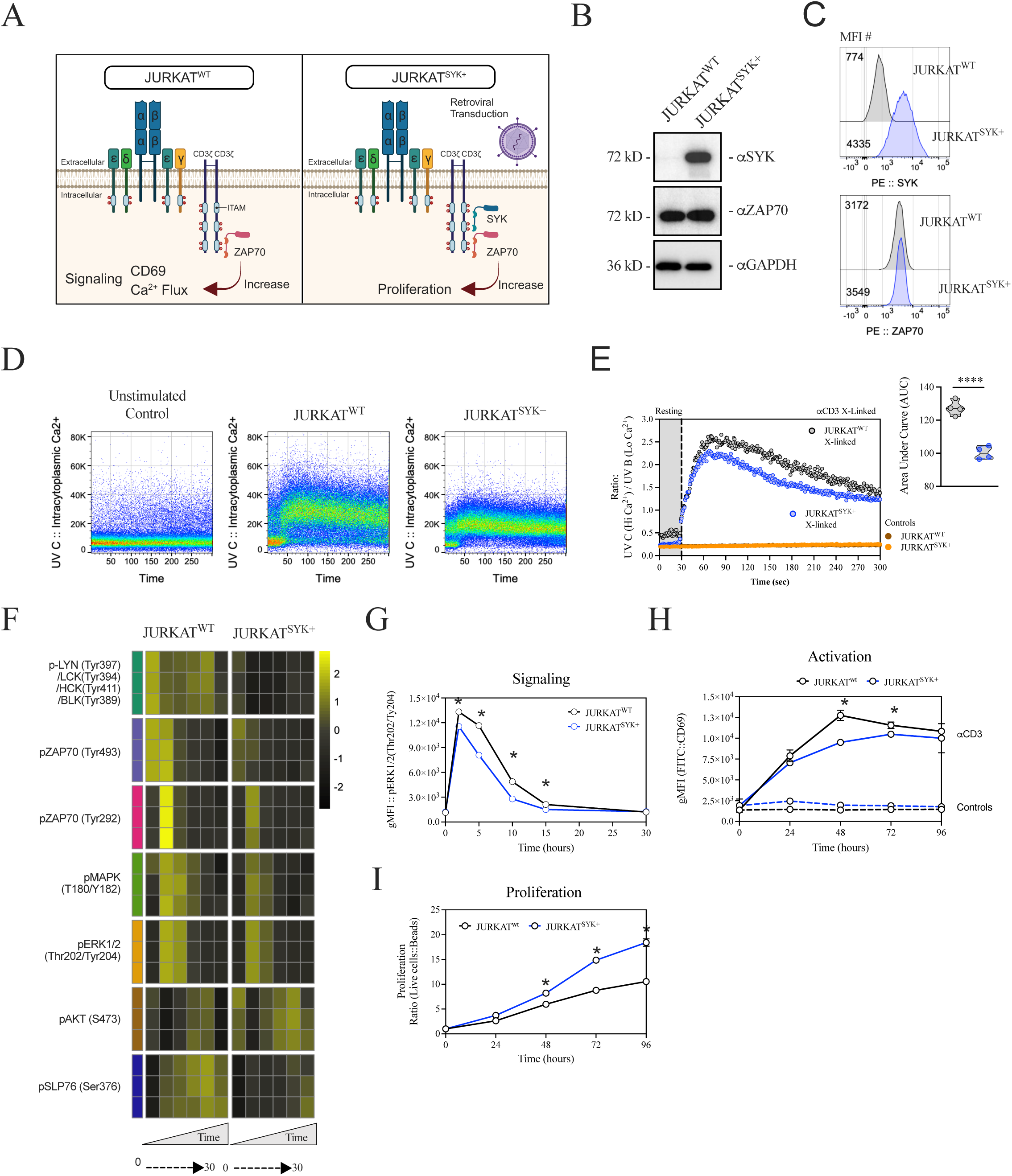
SYK expression in Jurkat T cells diminishes ITAM signaling while promoting proliferation. **(A-H)** Human T cell line Jurkat, constitutively expressing endogenous ZAP70, was transduced with a retroviral vector expressing SYK. **(A)** Diagram of WT and SYK-expressing Jurkat cells with associated biological outcomes. **(B)** Western blot analysis of Jurkat cells with anti-SYK, anti-ZAP70, and loading control anti-GAPDH antibodies. **(C)** Intracellular SYK and ZAP70 were analyzed by flow cytometry. **(D)** Calcium influx assay of unstimulated (left graph), and anti-CD3 cross-linked activation Jurkat cells (right two graphs). **(E)** Summary of the calcium influx for WT and SYK-expressing Jurkat cells; far right graph represents the calculated area under the curve (AUC) for each cell line and replicates. **(F)** Heatmap of phosphorylation kinetics of anti-CD3 cross-linked Jurkat cells for each signaling molecule. **(G)** Representative line graphs of geometric mean fluorescent intensity or pERK1/2 (Thr202/Ty204) from data in F. **(H)** Plate-bound anti-CD3 stimulated Jurkat cells analyzed for CD69 gMFI for 96 hours. **(I)** Flow cytometric analysis of proliferation of Jurkat cells cultured with media and 200 U/ml IL2 as measured with counting beads over 96 hours. Sample t test with significance is shown as ****, P <0.0001. Data in E-I represent 3-4 biological replicates for two or more independent experiments.

To identify the transcriptional effects of SYK and ZAP70 in the KHYG1 and Jurkat cells, RNA-Seq was performed on WT, ΔSYK, and ΔZAP70 KHYG1 cells and WT and SYK-bearing Jurkat cells. At the basal state without deliberate ITAM-based receptor stimulation many genes Analysis (PCA) show that the transcriptional profile of ΔSYK KHYG1 were most similar to WT were upregulated and downregulated in the variant *ZAP70* and *SYK* cells. Principal Component KHYG1 cells (PC2; 20% variance explained), while ΔZAP70 KHYG1 cells were most variable (DE) genes were observed in ΔZAP70 KHYG1 cells compared to ΔSYK KHYG1 cells with 161 compared to the others (PC1; 64% variance explained) (Fig. 5A). More differentially expressed of 3148 total DE genes overlapping (Fig. 5B). Genetic pathway analysis revealed significantly enriched pathways in “protein tyrosine kinase activity”, “regulation of MAPK cascade”, “regulation of ERK1 and ERK2 cascade”, and “calcium ion binding” in ΔSYK KHYG1 cells compared to WT KHYG1 cells (left panel of Fig. 5C). ΔZAP70 KHYG1 cells revealed significantly enriched gene pathways in metabolism and growth compared to WT KHYG1 cells (right panel of Fig. 5C). Heatmap analysis of ΔZAP70 KHYG1 cells show enrichment of genes related to phosphatase activity, pyruvate metabolic, and glucose metabolic process compared to WT and ΔSYK KHYG1 cells (Fig. 5D). DE analysis of SYK-bearing Jurkat cells resulted in more significantly downregulated genes compared to WT Jurkat cells (Fig. 5E). Pathway analysis revealed significantly suppressed pathways in calcium-dependent “phospholipid binding”, “MAP kinase kinase activity”, “regulation of receptor signaling pathway via STAT”, and “activation of protein kinase activity” in SYK-bearing Jurkat cells compared to WT Jurkat cells (Fig. 5F). Together, these data support a negative role for SYK in regard to phosphorylation and transcriptional programing in NK and T cells, whereas ZAP70 is essential for these functions.

**Figure 5:**
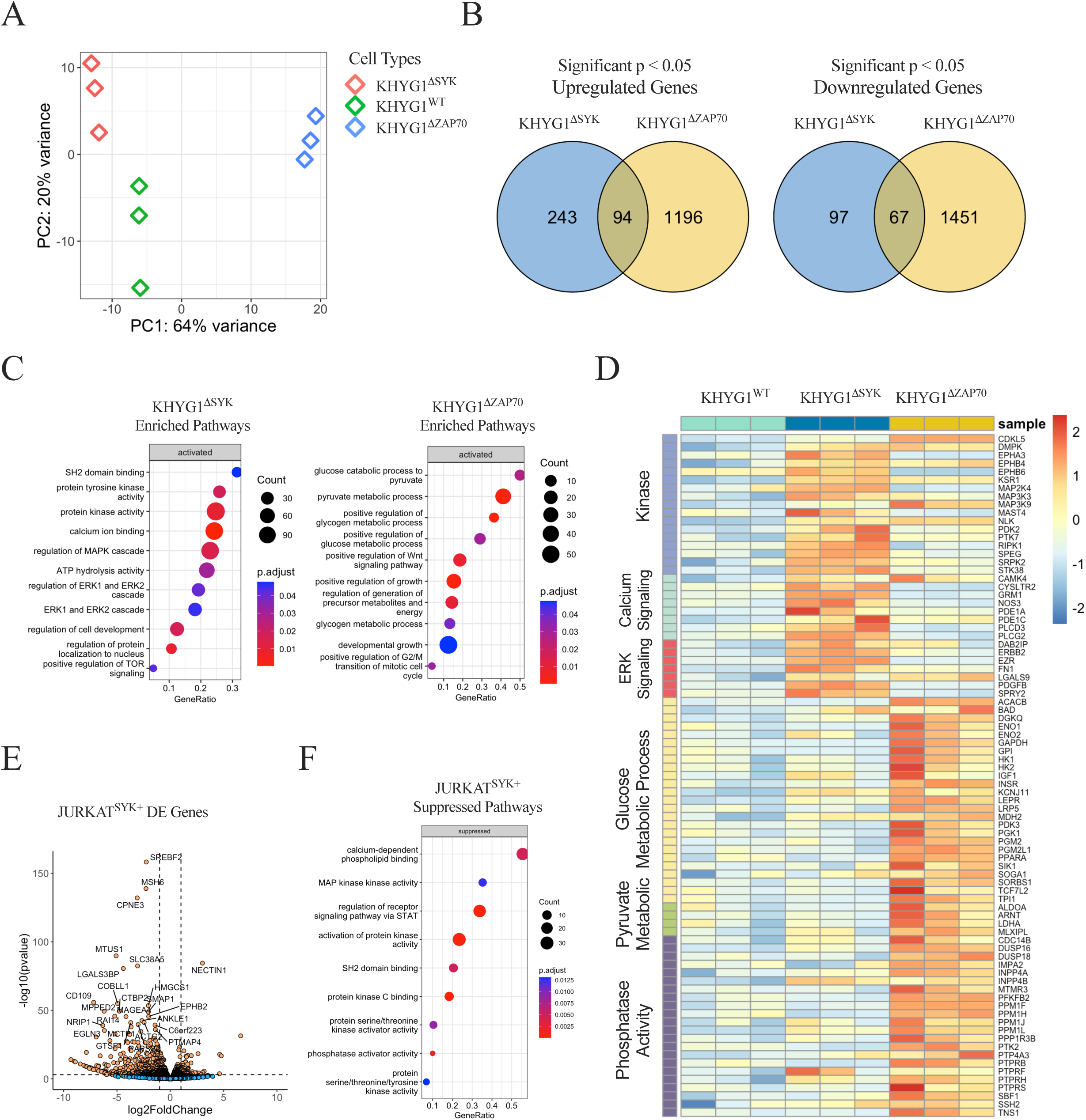
RNA sequencing of SYK and ZAP70 variant cell lines. **(A-F)** RNA sequencing of cells. **(A)** Principal component analysis (PCA) of normalized KHYG1 gene expression and percent variants on the axis. **(B)** Venn diagram of overlapping significant differential gene expression comparing WT KHYG1 with KHYG1 cells ablated of *SYK* or *ZAP70*. Top genes with significance of a p adjusted value of <0.05 were only considered in the Venn diagram. **(C)** Top-ranked gene set enrichment analysis of the top 10-11 chosen activated pathways enriched in KHYG1 cells depleted of *SYK* or *ZAP70*. **(D)** Heatmap of significant genes from the top pathways observed in C for each KHYG1 cell type. **(E)** Differential gene expression between WT and SYK-transduced Jurkat cells. Top significant genes were plotted, and orange dots represent significant genes; blue dots represent non-significant genes. **(F)** Top-ranked gene set enrichment analysis of the top 9 chosen suppressed pathways observed in Syk-transduced Jurkat cells. Data used for each figure consisted of 3 biological replicates for each cell line.

### Ablation of SYK in primary human CAR NK cells enhance tumor cell killing

The enhanced ITAM signaling in ΔSYK KHYG1 cells strongly suggest that ablation of *SYK* in primary human NK cells might enhance the function of CAR NK cells. Primary NK cells were transduced with transduced with a CD19-targeting CAR containing cytoplasmic signaling elements of CD28 and CD3ζ and then CRISPR editing was used to ablate *ZAP70* or *SYK* (Fig. 6A, B, C). We first confirmed by flow cytometry that genetic depletion of *SYK* or *ZAP70* did not alter surface expression of CD16, CD56, NKp30, NKp44, or NKp46 (data not shown). To test the influence of SYK and ZAP70, we conducted *in vitro* cytotoxicity assays with WT CAR NK cells, ΔZAP70 CAR NK cells, and ΔSYK CAR NK cells against SUPB15, a CD19-expressing B lymphoblast cell line, for 4 and 24 hour co-culture (Fig. 6D, E). *SYK*-ablated CAR NK cells mediated the most potent tumor-specific killing at 4 and 24 hours, whereas *ZAP70*-ablated CAR NK cells mediated the lowest tumor-specific killing compared to WT NK cells (Fig. 6D, E). Similarly, CAR NK cells ablated of *SYK* also show enhanced cytokine production (TNFα and IFNγ) and degranulation as measured by surface CD107a compared with WT CAR NK cells, whereas ΔZAP70 CAR NK cells had significantly diminished effector functions (Fig. 6F-I). Overall, our study shows that the ablation of *SYK* in NK cells enhanced CAR NK cell-mediated killing and increased production of effector cytokines against tumor targets (Fig. 6A).

**Figure 6:**
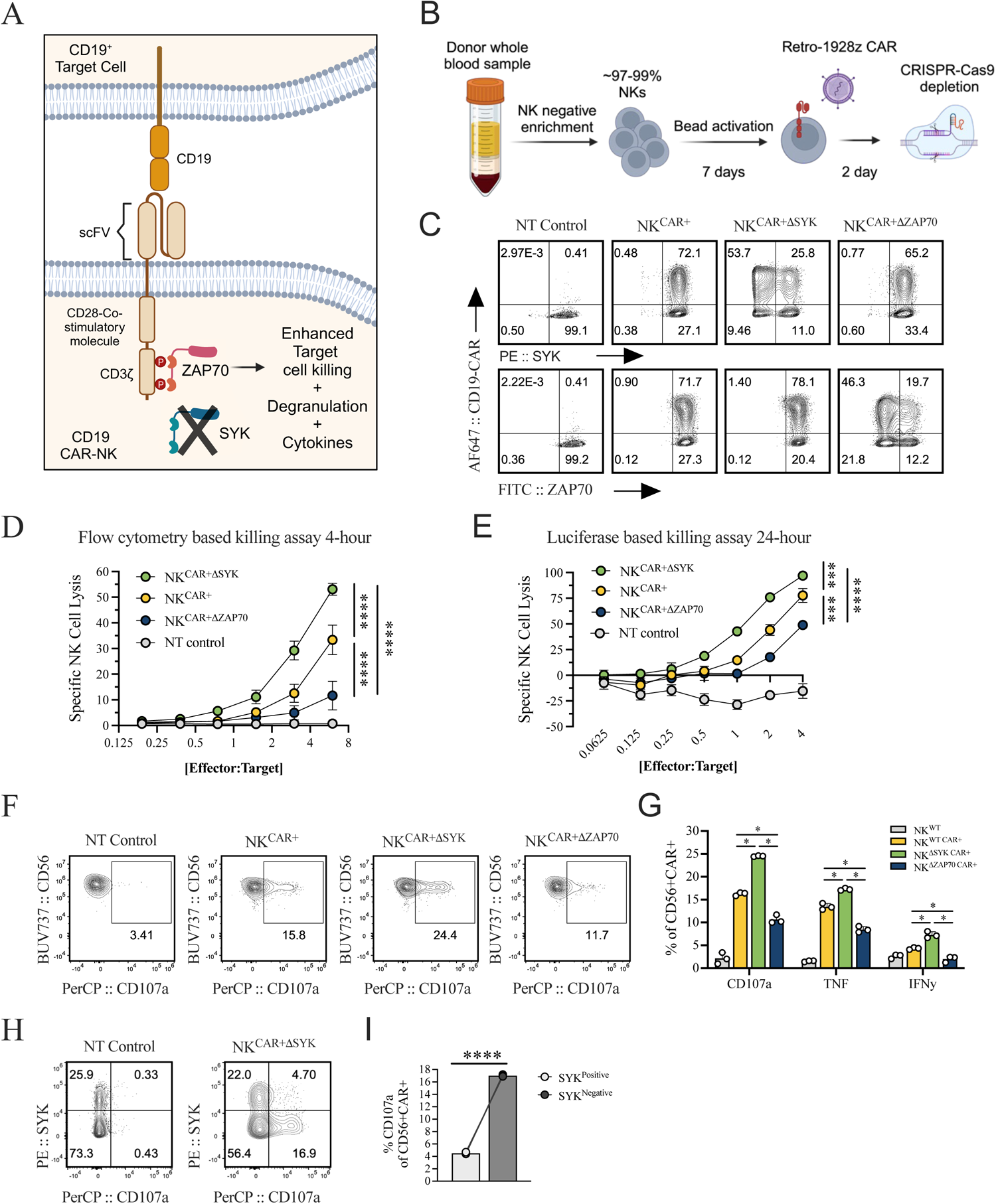
*SYK* ablation enhances CD19-CAR function in human primary NK cells. **(A-I)** *SYK*-ablated primary human NK cells demonstrated enhanced killing and cytokine responses against SUPB15 target cell line. **(A)** Depiction of ZAP70 signaling driving enhanced target cell killing, degranulation, and cytokine production in primary CAR NK cells. **(B)** Diagram of the 9-day introduction of the retroviral 1928ζ CAR construct into primary NK cells and subsequent depletion of *SYK* or *ZAP70* using CRISPR-Cas9. **(C)** Flow cytometry-based validation of CAR expression and *SYK* or *ZAP70* depletion in primary NK cells; gated on live CD3^−^CD14^−^CD19^−^CD56^+^ NK cells; non-transduced control (NT). **(D)** Summary graph of the 4-hour flow cytometry-based killing assay. **(E)** Summary graph of the 24-hour luciferase-based killing assay. **(F-I)** Markers identified after 4-hour co-culture with SUPB15 by flow cytometry. **(F)** Percentage of CD107a, IFNγ, and TNFα; plots of cells gated on live CD19-CAR^+^ NK cells. **(G)** Summary graph from data in F. **(H)** Flow cytometry plot of CD107a frequency, gated on live CD56^+^ CAR^+^ NK cells. **(I)** Summary plot of data shown in H. Data used in this figure consisted of 3 technical and 2 biological replicates per figure. Sample t test with significance is shown as ****, P <0.0001, ***, P <0.001, *, P <0.05.

## DISCUSSION

SYK and ZAP70 tyrosine kinases are present in all jawed vertebrates and are involved in immune receptor functions (11, 12). They contain SH2-binding and kinase domains that mediate their activity. SYK is more broadly expressed, being present in myeloid cells, B cells, T cells, and NK cells, whereas ZAP70 is predominantly expressed in T cells and NK cells. Mouse immature “double-negative” (DN) T cells in the thymus co-express both SYK and ZAP70, but typically lose SYK expression during maturation, although SYK is present in some γδ T cells, MAIT cells, and iNKT cells (www.immgen.org). Mice lacking *Zap70* are blocked at the “double-positive” stage of development and are largely devoid of mature αβ TCR T cells in peripheral lymphoid organs (11, 43). SYK is involved in pre-TCR signaling in DN thymocytes in mice and lack of Syk results in impaired T cell maturation (43). In humans, immature T cells in the thymus co-express SYK and ZAP70; however, unlike in mice humans with loss-of-function *ZAP70* genes lack CD8^+^ αβ TCR T cells in the periphery but have abundant CD4^+^ αβ TCR T cells due to expression and redundant function of SYK, albeit with defective function (44).

Mouse NK cells in all stages of development co-express SYK and ZAP70, but interestingly ILC2 and ILC3 lack SYK, but express ZAP70 (www.immgen.org). Normal numbers of NK cells are present in bone marrow chimeric *Rag2^−/−^Il2rg^−/−^* mice reconstituted with bone marrow with disrupted *Syk* genes and found that NK cell development was largely normal, potentially due to the redundant function of Zap70 (45). Natural cytotoxic activity was intact in *Syk^−/−^* mice, although they demonstrated slightly reduced ADCC activity (45). Mixed bone marrow chimeric *Rag2^−/−^Il2rg^−/−^*mice reconstituted with *Syk^−/−^* fetal liver cells demonstrated normal numbers of NK cells and natural cytotoxicity was intact, whereas B and T cells were absent (46). However, induction of IFNγ secretion by stimulation with agonist antibodies to CD16 (signaling through FcεR1γ) and Ly49D (signaling through DAP12) were absent, although surprisingly stimulation through NK1.1 was retained, implying a SYK and ZAP70-independent mechanism (46).

After clonal expansion induced by CMV infection human NK cells can acquire an adaptive or memory state characterized by the loss of SYK and FcεR1γ expression, accompanied by enhanced ADCC function (3, 4, 7). Human adaptive NK cells rely on CD3ζ and ZAP70 to activate these NK cells when initiated by the CD16 Fc receptor, similar to T cells using CD3ζ and ZAP70 to transmit TCR-induced signals. Loss of SYK and FcεR1γ does not occur in adaptive or memory mouse NK cells driven by mouse CMV infection. Unlike human CD3ζ, mouse CD3ζ cannot associate with CD16 so loss of FcεR1γ in mice results in total loss of ADCC function (15). Humans with loss-of-function *ZAP70* mutations have normal numbers of NK cells (44).

Given the loss of SYK in human adaptive NK cells, we investigated the roles of SYK and ZAP70 on downstream signal transduction in a model system ablating *SYK* or *ZAP70* in a clonal human NK cell line. Our study confirms and extends the recent findings of Dahlvang and colleagues (13), which demonstrated enhanced cytotoxicity in NK cells by ablation of *SYK*. Using a mechanistic *in silico* model, we showed the competition between ZAP70 and SYK in accessing a limited number of ITAMs leads to decrease in Ca^2+^ flux in WT compared to the ΔSYK KHYG1 NK cells. Our model predicts the presence of a higher number of ITAMs reduces the competition between ZAP70 and SYK for accessing phosphorylated ITAMs and thus can produce higher Ca^2+^ flux and potentially increased cytotoxicity in WT compared to ΔSYK NK cells. Our studies reveal that whereas SYK preferentially promotes proliferation, ZAP70 enhances Ca^2+^ influx and downstream activation when ITAM-based receptors are engaged. Similar results were obtained when Jurkat T cells endogenously expressing ZAP70 were transduced with SYK; proliferation was enhanced but TCR-induced activation was diminished. Analysis of KHYG1 NK cells and Jurkat cells expressing ZAP70 and/or SYK revealed extensive transcriptional alterations impacting pathways involved in MAPK, ERK1, and ERK2, Ca^2+^, metabolism, and phosphatase activity. Ablation of *ZAP70* or *SYK* in primary NK cells expressing a CD19-targeted CAR confirmed superior effector function in NK cell lacking SYK using *in vitro* assays to measure cytotoxicity and cytokine production. Further studies will be needed to evaluate the function of these *SYK*-ablated CAR NK cells *in vivo* and effects on proliferation and durability. Collectively, these studies reveal differential roles for SYK and ZAP70 in T and NK cells in regard to ITAM-mediated signaling, proliferation, and effector function.

## Acknowledgements

We thank Michael Blinov for introducing rule-based model in the Virtual Cell software. We thank Dr. James R. Faeder and Dr. Ali Sinan for assisting in technical questions related to PyBioNetGen library. IN thanks to HPC facility Franklin at the Nationwide Children’s Hospital for providing computational resources.

## REFERENCES

1. Li, Y., K. Rezvani, and H. Rafei. 2023. Next-generation chimeric antigen receptors for T- and natural killer-cell therapies against cancer. Immunol Rev 320: 217–235.

2. Lanier, L. L. 2005. NK cell recognition. Annu Rev Immunol 23: 225–274.

3. Lee, J., T. Zhang, I. Hwang, A. Kim, L. Nitschke, M. Kim, J. M. Scott, Y. Kamimura, L. L. Lanier, and S. Kim. 2015. Epigenetic modification and antibody-dependent expansion of memory-like NK cells in human cytomegalovirus-infected individuals. Immunity 42: 431–442.

4. Schlums, H., F. Cichocki, B. Tesi, J. Theorell, V. Beziat, T. D. Holmes, H. Han, S. C. Chiang, B. Foley, K. Mattsson, S. Larsson, M. Schaffer, K. J. Malmberg, H. G. Ljunggren, J. S. Miller, and Y. T. Bryceson. 2015. Cytomegalovirus infection drives adaptive epigenetic diversification of NK cells with altered signaling and effector function. Immunity 42: 443–456.

5. Guma, M., A. Angulo, C. Vilches, N. Gomez-Lozano, N. Malats, and M. Lopez-Botet. 2004. Imprint of human cytomegalovirus infection on the NK cell receptor repertoire. Blood 104: 3664–3671.

6. Lopez-Verges, S., J. M. Milush, B. S. Schwartz, M. J. Pando, J. Jarjoura, V. A. York, J. P. Houchins, S. Miller, S. M. Kang, P. J. Norris, D. F. Nixon, and L. L. Lanier. 2011. Expansion of a unique CD57(+)NKG2Chi natural killer cell subset during acute human cytomegalovirus infection. Proc Natl Acad Sci U S A 108: 14725–14732.

7. Hwang, I., T. Zhang, J. M. Scott, A. R. Kim, T. Lee, T. Kakarla, A. Kim, J. B. Sunwoo, and S. Kim. 2012. Identification of human NK cells that are deficient for signaling adaptor FcRgamma and specialized for antibody-dependent immune functions. Int Immunol 24: 793–802.

8. Lanier, L. L., B. Corliss, J. Wu, and J. H. Phillips. 1998. Association of DAP12 with activating CD94/NKG2C NK cell receptors. Immunity 8: 693–701.

9. Lanier, L. L., B. C. Corliss, J. Wu, C. Leong, and J. H. Phillips. 1998. Immunoreceptor DAP12 bearing a tyrosine-based activation motif is involved in activating NK cells. Nature 391: 703–707.

10. Mariuzza, R. A., P. Agnihotri, and J. Orban. 2020. The structural basis of T-cell receptor (TCR) activation: An enduring enigma. J Biol Chem 295: 914–925.

11. Au-Yeung, B. B., N. H. Shah, L. Shen, and A. Weiss. 2018. ZAP-70 in Signaling, Biology, and Disease. Annu Rev Immunol 36: 127–156.

12. Mocsai, A., J. Ruland, and V. L. Tybulewicz. 2010. The SYK tyrosine kinase: a crucial player in diverse biological functions. Nat Rev Immunol 10: 387–402.

13. Dahlvang, J. D., J. K. Dick, J. A. Sangala, P. R. Kennedy, E. J. Pomeroy, K. M. Snyder, J. M. Moushon, C. E. Thefaine, J. Wu, S. E. Hamilton, M. Felices, J. S. Miller, B. Walcheck, B. R. Webber, B. S. Moriarity, and G. T. Hart. 2023. Ablation of SYK Kinase from Expanded Primary Human NK Cells via CRISPR/Cas9 Enhances Cytotoxicity and Cytokine Production. J Immunol 210: 1108–1122.

14. Huang, R. S., M. C. Lai, H. A. Shih, and S. Lin. 2021. A robust platform for expansion and genome editing of primary human natural killer cells. J Exp Med 218.

15. Aguilar, O. A., L. K. Fong, K. Ishiyama, W. F. DeGrado, and L. L. Lanier. 2022. The CD3zeta adaptor structure determines functional differences between human and mouse CD16 Fc receptor signaling. J Exp Med 219.

16. Dobin, A., C. A. Davis, F. Schlesinger, J. Drenkow, C. Zaleski, S. Jha, P. Batut, M. Chaisson, and T. R. Gingeras. 2013. STAR: ultrafast universal RNA-seq aligner. Bioinformatics 29: 15–21.

17. Love, M. I., W. Huber, and S. Anders. 2014. Moderated estimation of fold change and dispersion for RNA-seq data with DESeq2. Genome Biol 15: 550.

18. Schaff, J., C. C. Fink, B. Slepchenko, J. H. Carson, and L. M. Loew. 1997. A general computational framework for modeling cellular structure and function. Biophysical journal 73: 1135–1146.

19. Faeder, J. R., M. L. Blinov, and W. S. Hlavacek. 2009. Rule-based modeling of biochemical systems with BioNetGen. Systems biology: 113–167.

20. Blinov, M. L., J. C. Schaff, D. Vasilescu, I. I. Moraru, J. E. Bloom, and L. M. Loew. 2017. Compartmental and spatial rule-based modeling with virtual cell. Biophysical journal 113: 1365–1372.

21. Leibson, P. J. 1997. Signal transduction during natural killer cell activation: inside the mind of a killer. Immunity 6: 655–661.

22. Matalon, O., and M. Barda-Saad. 2016. Cbl ubiquitin ligases mediate the inhibition of natural killer cell activity. Communicative & integrative biology 9: e1216739.

23. Lanier, L. L., G. Yu, and J. H. Phillips. 1989. Co-association of CD3ζ with a receptor (CD16) for IgG Fc on human natural killer cells. Nature 342: 803–805.

24. Lanier, L. L. 2003. Natural killer cell receptor signaling. Current opinion in immunology 15: 308–314.

25. Wu, Z., S. Park, C. M. Lau, Y. Zhong, S. Sheppard, J. C. Sun, J. Das, G. Altan-Bonnet, and K. C. Hsu. 2021. Dynamic variability in SHP-1 abundance determines natural killer cell responsiveness. Science signaling 14: eabe5380.

26. Wang, H.-Y., Y. Altman, D. Fang, C. Elly, Y. Dai, Y. Shao, and Y.-C. Liu. 2001. Cbl promotes ubiquitination of the T cell receptor ζ through an adaptor function of Zap-70. Journal of Biological Chemistry 276: 26004–26011.

27. Duan, L., A. L. Reddi, A. Ghosh, M. Dimri, and H. Band. 2004. The Cbl family and other ubiquitin ligases: destructive forces in control of antigen receptor signaling. Immunity 21: 7–17.

28. Au Yeung, B. B., S. Deindl, L. Y. Hsu, E. H. Palacios, S. E. Levin, J. Kuriyan, and A. Weiss. 2009. The structure, regulation, and function of ZAP 70. Immunological reviews 228: 41–57.

29. Shah, N. H., Q. Wang, Q. Yan, D. Karandur, T. A. Kadlecek, I. R. Fallahee, W. P. Russ, R. Ranganathan, A. Weiss, and J. Kuriyan. 2016. An electrostatic selection mechanism controls sequential kinase signaling downstream of the T cell receptor. Elife 5: e20105.

30. Mukherjee, S., J. Zhu, J. Zikherman, R. Parameswaran, T. A. Kadlecek, Q. Wang, B. Au-Yeung, H. Ploegh, J. Kuriyan, and J. Das. 2013. Monovalent and multivalent ligation of the B cell receptor exhibit differential dependence upon Syk and Src family kinases. Science signaling 6: ra1-ra1.

31. Sneddon, M. W., J. R. Faeder, and T. Emonet. 2011. Efficient modeling, simulation and coarse-graining of biological complexity with NFsim. Nature methods 8: 177–183.

32. Atri, A., J. Amundson, D. Clapham, and J. Sneyd. 1993. A single-pool model for intracellular calcium oscillations and waves in the Xenopus laevis oocyte. Biophysical Journal 65: 1727–1739.

33. Parys, J. B., S. W. Sernett, S. DeLisle, P. M. Snyder, M. J. Welsh, and K. P. Campbell. 1992. Isolation, characterization, and localization of the inositol 1, 4, 5-trisphosphate receptor protein in Xenopus laevis oocytes. Journal of Biological Chemistry 267: 18776–18782.

34. Ali Sinan Salgam, J. R. F. 2021. PyBioNetGen - A lightweight BioNetGen CLI.

35. Archdeacon, T. J. 1994. Correlation and regression analysis: a historian’s guide. Univ of Wisconsin Press.

36. Steel, R. G. D., and J. H. Torrie. 1960. Principles and procedures of statistics. Principles and procedures of statistics.

37. Plas, D. R., R. Johnson, J. T. Pingel, R. J. Matthews, M. Dalton, G. Roy, A. C. Chan, and M. L. Thomas. 1996. Direct regulation of ZAP-70 by SHP-1 in T cell antigen receptor signaling. Science 272: 1173–1176.

38. Bu, J. Y., A. S. Shaw, and A. C. Chan. 1995. Analysis of the interaction of ZAP-70 and syk protein-tyrosine kinases with the T-cell antigen receptor by plasmon resonance. Proceedings of the National Academy of Sciences 92: 5106–5110.

39. Zhao, Q., and A. Weiss. 1996. Enhancement of lymphocyte responsiveness by a gain-of-function mutation of ZAP-70. Mol Cell Biol 16: 6765–6774.

40. Chu, D. H., N. S. van Oers, M. Malissen, J. Harris, M. Elder, and A. Weiss. 1999. Pre-T cell receptor signals are responsible for the down-regulation of Syk protein tyrosine kinase expression. J Immunol 163: 2610–2620.

41. Krishnan, S., V. G. Warke, M. P. Nambiar, G. C. Tsokos, and D. L. Farber. 2003. The FcR gamma subunit and Syk kinase replace the CD3 zeta-chain and ZAP-70 kinase in the TCR signaling complex of human effector CD4 T cells. J Immunol 170: 4189–4195.

42. Abraham, R. T., and A. Weiss. 2004. Jurkat T cells and development of the T-cell receptor signalling paradigm. Nat Rev Immunol 4: 301–308.

43. Palacios, E. H., and A. Weiss. 2007. Distinct roles for Syk and ZAP-70 during early thymocyte development. J Exp Med 204: 1703–1715.

44. Elder, M. E., D. Lin, J. Clever, A. C. Chan, T. J. Hope, A. Weiss, and T. G. Parslow. 1994. Human severe combined immunodeficiency due to a defect in ZAP-70, a T cell tyrosine kinase. Science 264: 1596–1599.

45. Colucci, F., M. Turner, E. Schweighoffer, D. Guy-Grand, V. Di Bartolo, M. Salcedo, V. L. Tybulewicz, and J. P. Di Santo. 1999. Redundant role of the Syk protein tyrosine kinase in mouse NK cell differentiation. J Immunol 163: 1769–1774.

46. Colucci, F., E. Schweighoffer, E. Tomasello, M. Turner, J. R. Ortaldo, E. Vivier, V. L. Tybulewicz, and J. P. Di Santo. 2002. Natural cytotoxicity uncoupled from the Syk and ZAP-70 intracellular kinases. Nat Immunol 3: 288–294.

