## Supplementary materials for "Spleen Tyrosine Kinase (SYK) negatively regulates ITAM-mediated human NK cell signaling and CD19-CAR NK cell efficacy"

Supplementary Figure 1

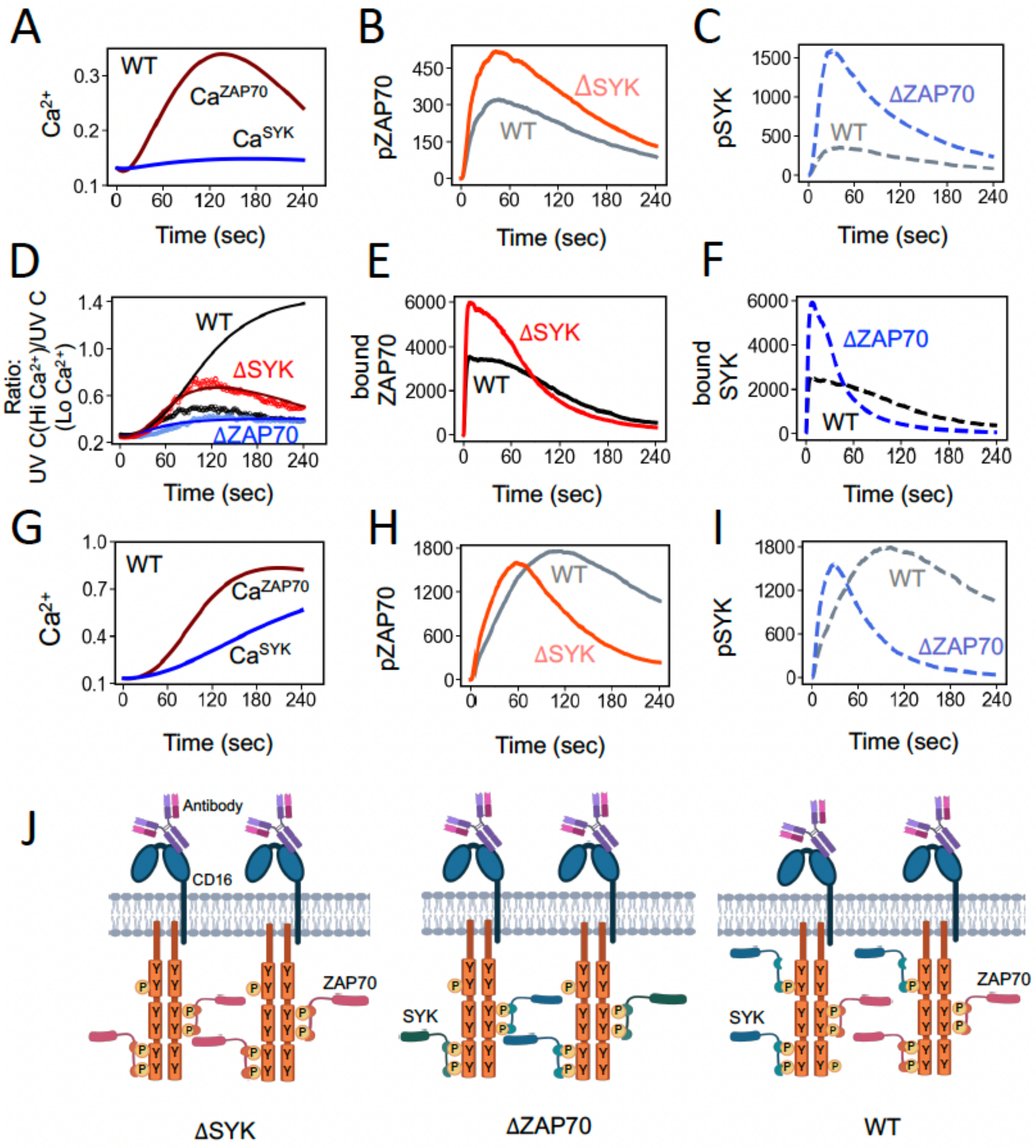


K

| ITAM access to ZAP70 | | ITAM access to SYK | |
|---|---|---|---|
| ΔSYK | WT | ΔZAP70 | WT |
| ++++ | +++ | ++++ | +++ |

ITAM number high

L

| ITAM access to ZAP70 | | ITAM access to SYK | |
|---|---|---|---|
| ΔSYK | WT | ΔZAP70 | WT |
| +++++ | +++++ | +++++ | +++++ |

Limiting case: ITAM number very high

**Figure S1: (A-C) Availability of ITAMs for ZAP70 and SYK at low concentrations (80 per μm$^2$) of CD16 and CD3ζ- homodimers.** (A) $Ca^{++}$ flux in WT shows the contributions of $Ca^{++}$ from ZAP70 and SYK, respectively. (B) Comparison of number of phospho-ZAP70 is shown between ΔSYK (solid orange line) and WT (solid gray line). (C) Comparison of number of phospho-SYK is shown between ΔZAP70 (dashed blue line) and WT (dashed gray line). (B-C) are generated within the simulation box proximal to the cell membrane. (D-I) **Availability of ITAMs for ZAP70 and SYK at high concentrations (200 per μm$^2$) of CD16 and CD3ζ- homodimers.** (D) *In silico* model is trained by fitting $Ca^{++}$ for ΔSYK and ΔZAP70 NK cells. Prediction of $Ca^{++}$ flux in WT using *in silico* model is shown in black solid line, which is higher than ΔSYK cells. (E) Comparison in number of ITAM-bound ZAP70 is shown between ΔSYK (solid red) and WT (solid black). (F) Comparison in number of ITAM-bound SYK is shown between ΔZAP70 (dashed blue) and WT (dashed black). (G) $Ca^{++}$ flux in WT shows the contributions of $Ca^{++}$ from ZAP70 and SYK, respectively. (H) Comparison in number of phospho-ZAP70 is shown between ΔSYK (solid orange line) and WT (solid gray line). (I) Comparison in number of phospho-SYK is shown between ΔZAP70 (dashed royal blue line) and WT (dashed gray line). (E-F) and (H-I) are generated within the simulation box proximal to the cell membrane. **(J-L) Mechanism of ITAM accessibility by ZAP70 and SYK for higher ITAM number.** (J) ZAP70 and SYK have more access to ITAMs for higher CD16 and CD3ζ homodimer concentrations in ΔSYK, ΔZAP70, and WT cells, which decreases the competition between ZAP70 and SYK for accessing phosphorylated ITAMs in the WT. (K) and (L) summarize the decrease in the competition to bind to ITAMs when the numbers of ITAMs are higher or substantially higher compared to the case investigated in Fig. 2H in the main text.

### Supplementary Table 1

| **A. Parameter values used in simulation** | | | |
|---|---|---|---|
| **Species concentration** | **Values used in Simulation** | **Source** | **Remarks** |
| Ligand (Anti-CD16 antibody) | 60 molecules/$\mu m^3$ | Estimated | We assumed the antibody molecules in a volume of 1.0 μm × 25 $\mu m^2$ (area of the plasma membrane in the simulation box) bind to the CD16 molecules in the simulation. |
| CD16 Receptor | 80 molecules /$\mu m^2$ | Fig. 1B in (1) | Ref. (1) estimates 70000-110000 CD16 receptor per NK cells obtained from PBMCs. Assuming the diameter of a primary human NK cell to be ~7 μm, the concentration of CD16 in the plasma membrane is ~ 450 molecules /$\mu m^2$. For KHYG1 cell line we have assumed a smaller concentration of CD16 than the primary NK cells in the PBMCs. |
| CD3ζ | 80 molecules/$\mu m^2$ | Assumption | Surface expressions of human CD16 is stabilized by CD3ζ adaptors (Fig. S3 in (2)). We assumed the concentrations of CD16 are CD3ζ are similar. |
| LCK | 390 molecules /$\mu m^2$ | Table S1 in (3) | Taken from the values measured for Jurkat T cells. |
| SHP-1 | 0.3 μM =180 molecules/$\mu m^3$ | Table S1 in (3) | Value used using estimation of SHP-2 in Jurkat T cells, maxQB database (4) |
| Cbl | 141 molecules/$\mu m^3$ | See Table S2 | Estimated by fitting our model to the $Ca^{++}$ kinetics data. |

| **Kinetic reaction rates** | **Values used in Simulation** | **Region of reaction** | **Source** | **Remarks** |
|---|---|---|---|---|

| | | | | |
|---|---|---|---|---|
| CD16 binding with anti-CD16 antibody ($k_{ON}$) | $1.7 \times 10^{-5}$ µ$m^3$ $s^{-1}$ molecule $^{-1}$ | Extracellular volume ($V_e$)<br><br>$V_e$ = 5µm × 5µm × 0.002µm = 0.05 µ$m^3$ | | CD16 are crosslinked by anti-CD16 antibody (2). We considered here, $K_D$ to be ~ 1 µM. |
| CD16 unbinding from anti-CD16 antibody ($k_{OFF}$) | $9.95 \times 10^{-3}$ $s^{-1}$≅ 0.01 $s^{-1}$ | Extracellular volume ($V_e$) | Table 1 in (5) | Value corresponds to IgG1 Fc (WT) unbinding from human CD16A – 158V in ref. (5). |
| CD3ζ adaptor (homodimer) binding to the CD16 receptor | $2{\times}10^{-3}$ µ$m^2$ $s^{-1}$ molecule$^{-1}$ | Membrane (A)<br>A=<br>5µm × 5µm = 25 µ$m^2$ | Assumption | |
| CD3ζ adaptor (homodimer) unbinding from CD16 receptor | 0.01 $s^{-1}$ | Membrane (A) | Assumption | |
| Phosphorylation of CD3ζ ITAMs by LCK | $1.5 \times 10^{-2}$ µ$m^2$ $s^{-1}$ | Membrane (A) | Table 1 in (6) | Values of $k_{cat}/k_M$ for Lck pY394-pY505 were calculated by fitting Michaelis Menten model in ref. (6). |
| Dephosphorylation of ITAMs on CD3ζ | 0.001 $s^{-1}$ | Membrane (A) | | Assumption |
| ZAP70 binding to fully phosphorylated ITAM | 5 x $10^6$ $M^{-1}s^{-1}$ =5 (µM s)$^{-1}$=$8.3 \times 10^{-3}$ µ$m^3$ $s^{-1}$ molecule$^{-1}$ | Cytosolic volume ($V_c$)<br>$V_c$ = 5 µm× 5µm×1 µm = 25 µ$m^3$ | (7) | Measured values reported in ref. (7) for T cells.<br><br>$K_a$ (association rate) |

| | | | | |
|---|---|---|---|---|
| | | | | = 5 x $10^6$ $M^{-1}s^{-1}$ |
| ZAP70 unbinding from fully phosphorylated ITAM | 0.125 $s^{-1}$ | Cytosolic volume ($V_c$) | (7) | Measured values reported in ref. (7) for T cells. $K_a$ (association rate)= 5 x $10^6$ $M^{-1}s^{-1}$, and, $K_D$=25 nM We approximated the unbinding rate as $K_a \times K_D$ using the values reported in ref. (7) . |
| ZAP70 phosphorylation by LCK | 3 ×$10^{-5}$ $(nMs)^{-1}$=0.03 $(\mu M\ s)^{-1}$ = 4.98 × $10^{-5}$ $\mu m^3\ s^{-1}$ molecule $^{-1}$ | Cytosolic volume ($V_c$) | Table 2 in (8) | Value used for phosphorylation of ZAP70 Y493 by Lck. |
| ZAP70 (free or bound) dephosphorylation by SHP-1 | 0.034 $(\mu M\ s)^{-1}$ = 5.64 × $10^{-5}$ $\mu m^3\ s^{-1}$ molecule $^{-1}$ | Cytosolic volume ($V_c$) | Table 2 in (9) | Catalytic rate of SHP-1(tethered) for substrate PEG12 calculated in (9). |
| SYK binding to fully phosphorylated ITAM | 5 $(\mu M\ s)^{-1}$=8.3 × $10^{-3}$ $\mu m^3\ s^{-1}$ molecule $^{-1}$ | Cytosolic volume ($V_c$) | (7) | SYK and ZAP70 affinities to fully phosphorylated ITAMs correspond to $K_a$ (association rate)= 5 x $10^6$ $M^{-1}s^{-1}$ reported in (7). |
| SYK binding to partially phosphorylated ITAM | 8.3 × $10^{-4}$ $\mu m^3\ s^{-1}$ molecule | Cytosolic volume ($V_c$) | Assumption | We assumed SYK binds to the partially phosphorylated |

| | | | | |
|---|---|---|---|---|
| | | | | state of the ITAM (state U) with a 10 × lower rate (=250 nM) |
| SYK unbinding from fully or partially phosphorylated ITAM | 0.125 $s^{-1}$ | Cytosolic volume ($V_c$) | (7) | SYK and ZAP70 affinities to the ITAM correspond to $K_a$ (association rate)= 5 x $10^6$ $M^{-1}s^{-1}$ and $K_D$=25 nM reported in ref. (7). |
| Catalytically active SYK phosphorylating ITAM | 1.21 $s^{-1}$ | Cytosolic volume ($V_c$) | Table S1 in (10) | Assumed following BCR signaling where phospho-ITAM bound catalytically active SYK fully phosphorylates the ITAM it is attached to (11). |
| Basally active SYK phosphorylating ITAM | 8.4 $\mu M^{-1}s^{-1}$ =0.014 $\mu m^3$ $s^{-1}$ molecule $^{-1}$ | Cytosolic volume ($V_c$) | Table S1 in (10) | Assumed following BCR signaling where phospho-ITAM bound basally active SYK fully phosphorylates other ITAMs (12). |
| Catalytically active SYK phosphorylating ITAM | 15.1 $\mu M^{-1}s^{-1}$=0.025 $\mu m^3$ $s^{-1}$ molecule $^{-1}$ | Cytosolic volume ($V_c$) | Table S1 in (10) | Assumed following BCR signaling where phospho-ITAM bound catalytically active SYK fully phosphorylates other ITAMs. |

| | | | | |
|---|---|---|---|---|
| SYK or ZAP70 (bound or free) dephosphorylation rate by phosphatase SHP-1 | $5.64 \times 10^{-5}$ μm³ s⁻¹ molecule⁻¹ | Cytosolic volume ($V_c$) | (13) | Assumed similar dephosphorylation rates for pSYK and pZAP70. |
| SHP-1 binding rate to fully phosphorylated ITAM | 0.25 $(\mu M\ s)^{-1}$ $= 4.15 \times 10^{-4}\ \mu m^3\ s^{-1}$ $molecule^{-1}$ | Cytosolic volume ($V_c$) | Table 2 in (9) SI in (14) | Value measured for PEG12 peptides in SPR experiment. SHP-1 injected over immobilized phosphorylated PEG-PD1 peptides. |
| SHP-1 unbinding rate from fully phosphorylated ITAM | 1.7 $s^{-1}$ | | Table 2 in (9) | Value measured for PEG12 peptides. |
| SHP-1 binding to partially phosphorylated ITAM | 0.25 $(\mu M\ s)^{-1}$ $= 4.15 \times 10^{-4}\ \mu m^3\ s^{-1}$ $molecule^{-1}$ | Cytosolic volume ($V_c$) | Table 2 in (9) SI in (14) | We considered the same rates for binding of SHP-1 to partially or fully phosphorylated ITAMs. |
| SHP-1 unbinding from partially phosphorylated ITAM | 1.7 $s^{-1}$ | | Table 2 in (9) | |
| Cbl binds to ITAM bound ZAP70 or SYK | 0.4 $(\mu M\ s)^{-1}$ $= 6.7 \times 10^{-4}\ \mu m^3\ s^{-1}$ $molecule^{-1}$ | | Assumption | |
| Cbl unbinds from ITAM bound ZAP70 or SYK | 0.004 $s^{-1}$ | | Assumption | |
| ITAM bound ZAP70 or SYK degradation rate by Cbl | 0.07 $s^{-1}$ | | Assumption | Cbl binds to the phospho-ITAM bound ZAP70 and |

| | | | | |
|---|---|---|---|---|
| | | | | SYK and degrades the ITAM bound molecule (ZAP70 or SYK) and whole CD16 (15). |
| Free pZAP70 or pSYK dephosphorylation | 0.01 $s^{-1}$ | | Assumption | Represents dephosphorylation of ZAP70 and SYK by other phosphatases than SHP-1. |
| Parameter $k_1$ in $Ca^{++}$ ODE | 0.7 μM | | (16, 17) | Determined by fitting data for Xenopus oocyte in ref. (17) |
| parameter $k_2$ in $Ca^{++}$ ODE | 0.7 μM | | (16, 17) | Determined by fitting data for Xenopus oocyte in ref.(17) |
| Parameter $b$ in $Ca^{++}$ ODE | 0.111 | | (16, 17) | Determined by fitting data for Xenopus oocyte in ref. (17) |
| Rate for the kinetic proof-reading reaction step | 0.01 $s^{-1}$ | | Assumption | |

**B: Estimated parameters from *in silico* model training for limited ITAMs (concentrations of CD16 and CD3ζ homodimers = 80 molecules/$\mu m^2$)**

| | **Parameters** | **Estimated values** |
|---|---|---|
| **CD16 signaling model** **CD16 signaling model** | initial ZAP70 concentration | 1131 molecules/$um^3$ |
| | initial SYK concentration | 754 molecules/$um^3$ |
| | SYK auto-phosphorylation rate | 0.0026 $s^{-1}$ |
| | SYK trans-phosphorylation rate | 0.0022 $\mu m^3$ $molecule^{-1}$ $s^{-1}$ |
| | Cbl concentration | 141 molecules/$\mu m^3$ |
| | **Parameters for ΔSYK** | **Parameters for ΔZAP70** |

| Ca$^{++}$ ODE model | Parameters | Estimated Values | Parameters | Estimated Values |
|---|---|---|---|---|
| | $C_{1Z}$ | 0.785 $s^{-1}$ | $C_{1S}$ | 0.058 $s^{-1}$ |
| | $C_{2Z}$ | 0.161 $\mu M^{-1}$ $s^{-1}$ | $C_{2S}$ | 0.0001 $\mu M^{-1}$ $s^{-1}$ |
| | $\gamma_Z$ | 0.012 $s^{-1}$ | $\gamma_S$ | 0.00103 $s^{-1}$ |

**C. Estimated parameters from *in silico* model training for higher ITAMs (concentrations of CD16 and CD3ζ homodimers = 200 per $\mu m^2$)**

| | Parameters | Estimated Values |
|---|---|---|
| **CD16 signaling model** | initial ZAP70 concentration | 896 molecules/$\mu m^3$ |
| | initial SYK concentration | 598 molecules/$\mu m^3$ |
| | SYK auto-phosphorylation rate | 0.014959 $s^{-1}$ |
| | SYK trans-phosphorylation rate | $2.823 \times 10^{-5}$ $\mu m^3$ $molecule^{-1}$ $s^{-1}$ |

| | Parameters for ZAP70 | | Parameters for SYK | |
|---|---|---|---|---|
| **Ca$^{++}$ ODE model** | **Parameters** | **Estimated Values** | **Parameters** | **Estimated Values** |
| | $C_{1Z}$ | 0.2243 $s^{-1}$ | $C_{1S}$ | 0.0545 $s^{-1}$ |
| | $C_{2Z}$ | 0.538418 $\mu M^{-1}$ $s^{-1}$ | $C_{2S}$ | 0.00107 $\mu M^{-1}$ $s^{-1}$ |
| | $\gamma_Z$ | 0.00559 $s^{-1}$ | $\gamma_S$ | 0.000264 $s^{-1}$ |

1. Hatjiharissi, E., L. Xu, D. D. Santos, Z. R. Hunter, B. T. Ciccarelli, S. Verselis, M. Modica, Y. Cao, R. J. Manning, and X. Leleu. 2007. Increased natural killer cell expression of CD16, augmented binding and ADCC activity to rituximab among individuals expressing the FcγRIIIa-158 V/V and V/F polymorphism. *Blood, the Journal of the American Society of Hematology* 110: 2561-2564.
2. Aguilar, O. A., L.-K. Fong, K. Ishiyama, W. F. DeGrado, and L. L. Lanier. 2022. The CD3ζ adaptor structure determines functional differences between human and mouse CD16 Fc receptor signaling. *Journal of Experimental Medicine* 219: e20220022.
3. Hui, E., J. Cheung, J. Zhu, X. Su, M. J. Taylor, H. A. Wallweber, D. K. Sasmal, J. Huang, J. M. Kim, and I. Mellman. 2017. T cell costimulatory receptor CD28 is a primary target for PD-1–mediated inhibition. *Science* 355: 1428-1433.
4. Schaab, C., T. Geiger, G. Stoehr, J. Cox, and M. Mann. 2012. Analysis of high accuracy, quantitative proteomics data in the MaxQB database. *Molecular & Cellular Proteomics* 11.
5. Ellwanger, K., U. Reusch, I. Fucek, S. Wingert, T. Ross, T. Müller, U. Schniegler-Mattox, T. Haneke, E. Rajkovic, and J. Koch. 2019. Redirected optimized cell killing (ROCK®): a highly versatile multispecific fit-for-purpose antibody platform for engaging innate immunity. In *MAbs*. Taylor & Francis. 899-918.
6. Hui, E., and R. D. Vale. 2014. In vitro membrane reconstitution of the T-cell receptor proximal signaling network. *Nature structural & molecular biology* 21: 133-142.
7. Bu, J. Y., A. S. Shaw, and A. C. Chan. 1995. Analysis of the interaction of ZAP-70 and syk protein-tyrosine kinases with the T-cell antigen receptor by plasmon resonance. *Proceedings of the National Academy of Sciences* 92: 5106-5110.
8. Arulraj, T., and D. Barik. 2018. Mathematical modeling identifies Lck as a potential mediator for PD-1 induced inhibition of early TCR signaling. *PloS one* 13: e0206232.
9. Clemens, L., M. Kutuzov, K. V. Bayer, J. Goyette, J. Allard, and O. Dushek. 2021. Determination of the molecular reach of the protein tyrosine phosphatase SHP-1. *Biophysical Journal* 120: 2054-2066.
10. Mukherjee, S., J. Zhu, J. Zikherman, R. Parameswaran, T. A. Kadlecek, Q. Wang, B. Au-Yeung, H. Ploegh, J. Kuriyan, and J. Das. 2013. Monovalent and multivalent ligation of the B cell receptor exhibit differential dependence upon Syk and Src family kinases. *Science signaling* 6: ra1-ra1.
11. Tsang, E., A. M. Giannetti, D. Shaw, M. Dinh, K. Joyce, S. Gandhi, H. Ho, S. Wang, E. Papp, and J. M. Bradshaw. 2008. Molecular mechanism of the Syk activation switch. *Journal of Biological Chemistry* 283: 32650-32659.
12. Rolli, V., M. Gallwitz, T. Wossning, A. Flemming, W. W. Schamel, C. Zürn, and M. Reth. 2002. Amplification of B cell antigen receptor signaling by a Syk/ITAM positive feedback loop. *Molecular cell* 10: 1057-1069.
13. Brumbaugh, K. M., B. A. Binstadt, D. D. Billadeau, R. A. Schoon, C. J. Dick, R. M. Ten, and P. J. Leibson. 1997. Functional role for syk tyrosine kinase in natural killer cell–mediated natural cytotoxicity. *The Journal of experimental medicine* 186: 1965-1974.
14. Das, J. 2010. Activation or tolerance of natural killer cells is modulated by ligand quality in a nonmonotonic manner. *Biophysical journal* 99: 2028-2037.
15. Matalon, O., and M. Barda-Saad. 2016. Cbl ubiquitin ligases mediate the inhibition of natural killer cell activity. *Communicative & integrative biology* 9: e1216739.

16. Atri, A., J. Amundson, D. Clapham, and J. Sneyd. 1993. A single-pool model for intracellular calcium oscillations and waves in the Xenopus laevis oocyte. *Biophysical Journal* 65: 1727-1739.
17. Parys, J. B., S. W. Sernett, S. DeLisle, P. M. Snyder, M. J. Welsh, and K. P. Campbell. 1992. Isolation, characterization, and localization of the inositol 1, 4, 5-trisphosphate receptor protein in Xenopus laevis oocytes. *Journal of Biological Chemistry* 267: 18776-18782.
